# Polyploid - diploid coexistence in the greater duckweed *Spirodela polyrhiza*

**DOI:** 10.1101/2025.01.07.631665

**Authors:** Frederik Mortier, Yves Van de Peer, Dries Bonte

## Abstract

Polyploidy, resulting from whole-genome duplication (WGD), is widespread among plants and originates in sympatry with their lower-ploidy progenitors. New polyploids can only succeed when they overcome the competitive disadvantage against their progenitors or through sufficient niche differentiation as resulting in a negative frequency dependent growth. Because polyploidy is frequently associated with cold, dry, and saline environments, stress is anticipated to be the key to polyploid success.

We examined the invasion of neotetraploid duckweed (*Spirodela polyrhiza*) strains in populations of their direct diploid progenitors, in control and salt stress conditions in replicated microcosms. We also tested the reverse scenario, that is, the invasion of diploids in neotetraploid populations, to determine if the initial proportion of tetraploids affects the outcome of competition.

Our results showed that the proportion of tetraploids declined in all tetraploid and all diploid invasions and it did so at a different rate than expected from only differences in intrinsic growth rate. Salt stress affected the decline in tetraploid proportion differently across strains. We also found evidence for negative frequency dependent growth that, nonetheless, was insufficient to overcome competitive disadvantages of neopolyploids towards their diploid progenitor.

We showcase a robust quantitative pipeline from flow cytometry of mixed-ploidy populations to population model fitting. In doing so, we demonstrate the important effect of competition and frequency dependency on neopolyploid establishment. Therefore, we caution for inferring neopolyploid success from intrinsic growth rates alone.

## Introduction

Polyploid organisms possess more than two sets of chromosomes as a consequence of whole genome duplication (WGD, Otto and Whitton, 2000; Stebbins, 1971, 1950). Unreduced gametes that are the result of cell cycle errors may pair with other unreduced gametes to form an autopolyploid in sympatry with its progenitors (Felber, 1991; Kauai et al., 2023; Ramsey and Schemske, 1998). The establishment of a polyploid variant is determined by the immediate phenotypic effect that accompanies such a drastic genomic change (Bomblies, 2020; Clo and Kolář, 2021), but remains to be fully understood to this day. A newly emerged polyploid, called neopolyploid, first needs to grow in numbers to establish a viable population while interacting with all co-occurring organisms. Barring a simultaneous colonization event, neopolyploids tend to emerge in sympatry, and in competition, with their abundant lower-ploidy progenitors. Therefore, mechanisms underlying polyploid establishment can only be unraveled by examining the outcome of competition between the neopolyploid and its progenitors.

Competition between reproductively isolated cytotypes, comparable to interspecific competition, may result in their coexistence or competitive exclusion based on the population growth rate of both cytotypes. However, many neopolyploids grow and reproduce slower than their direct progenitors (Clo and Kolář, 2021) and therefore are on first glance not expected to establish. The intrinsic growth rate is typically reduced in polyploids because of intrinsic costs associated with WGD, such as an increased polyploid cell size, larger N and P demands to produce DNA, and genomic shock (Anneberg et al., 2023; Comai, 2005; Guignard et al., 2016). Polyploid population growth also suffers from sexual incompatibility with their progenitor cytotype, which occurs at high frequency (Husband and Sabara, 2004; Ramsey and Schemske, 1998) leading to minority cytotype exclusion (MCE, Fowler and Levin, 1984; Husband, 2000; Levin, 1975; Rodríguez, 1996). These costs to polyploidy put a neopolyploid at an inherent disadvantage compared to its progenitors (Clo and Kolář, 2021).

While differences in intrinsic growth rates are important, the faith of two co-occurring cytotypes is also determined by their responses to intra- and intercytotype competition (Chesson, 2000; Mortier et al., 2024). A reduced intrinsic growth rate of the neopolyploid compared to its progenitor can be overcome if the polyploid copes better with competition (i.e., increasing population densities) across all cytotypes. A reduced intrinsic growth rate can also be overcome if the strength of intercytotype compared to intracytotype competition is reduced, which results in a negative effect of a cytotype’s frequency on its growth. Such negative frequency-dependency is considered a characteristic of niche differentiation and promotes species/cytotype coexistence (Fig. 1; Chase and Leibold, 2003; Gause, 1934). Alternatively, positive frequency-dependent effects on growth can cause a higher-frequency cytotype to maintain its dominance by the time a later cytotype arrives, so called priority effects (De Meester et al., 2016; Fukami, 2015). Examples of positive feedback are expected when, same-cytotype individuals facilitate each other’s growth or different-cytotype individuals interact more negatively than just resource competition (Fukami, 2015). Positive frequency-dependent effects counteract possible coexistence promoting effects of niche differentiation (Fowler and Levin, 1984; Grainger et al., 2019a). The outcome of competing populations, including effects of cytotype frequency, can be predicted by the mutual invasion criterion (Grainger et al., 2019b). The mutual invasion criterion poses that both populations can coexist stably (i.e., long-term) if both grow at low density when the competitor is at high density. Conversely, only one population may be able to invade the other but not *vice versa* (i.e., deterministic exclusion) or both populations may be unable to invade the other (i.e., priority effects, Fig. 1). Modern coexistence theory builds upon the mutual invasion criterion by quantifying the frequency-independent effects (relative fitness difference) and frequency-dependent effects (niche difference) on competition outcomes (Barabás et al., 2018; Chesson, 2000; Grainger et al., 2019a). Modern coexistence theory explains competition between cytotypes most effectively when cytotypes are (almost completely) reproductively isolated, such as is often the case with strong MCE, strong assortative mating and asexual reproduction (discussions and models on the effect of incomplete isolation: Gaynor et al., 2024; Irwin and Schluter, 2022). The expected survival or exclusion of a neopolyploid in competition with its progenitor is, therefore, determined by how polyploidy influences frequency independent and frequency dependent effects on population growth.

**Fig. 1:**
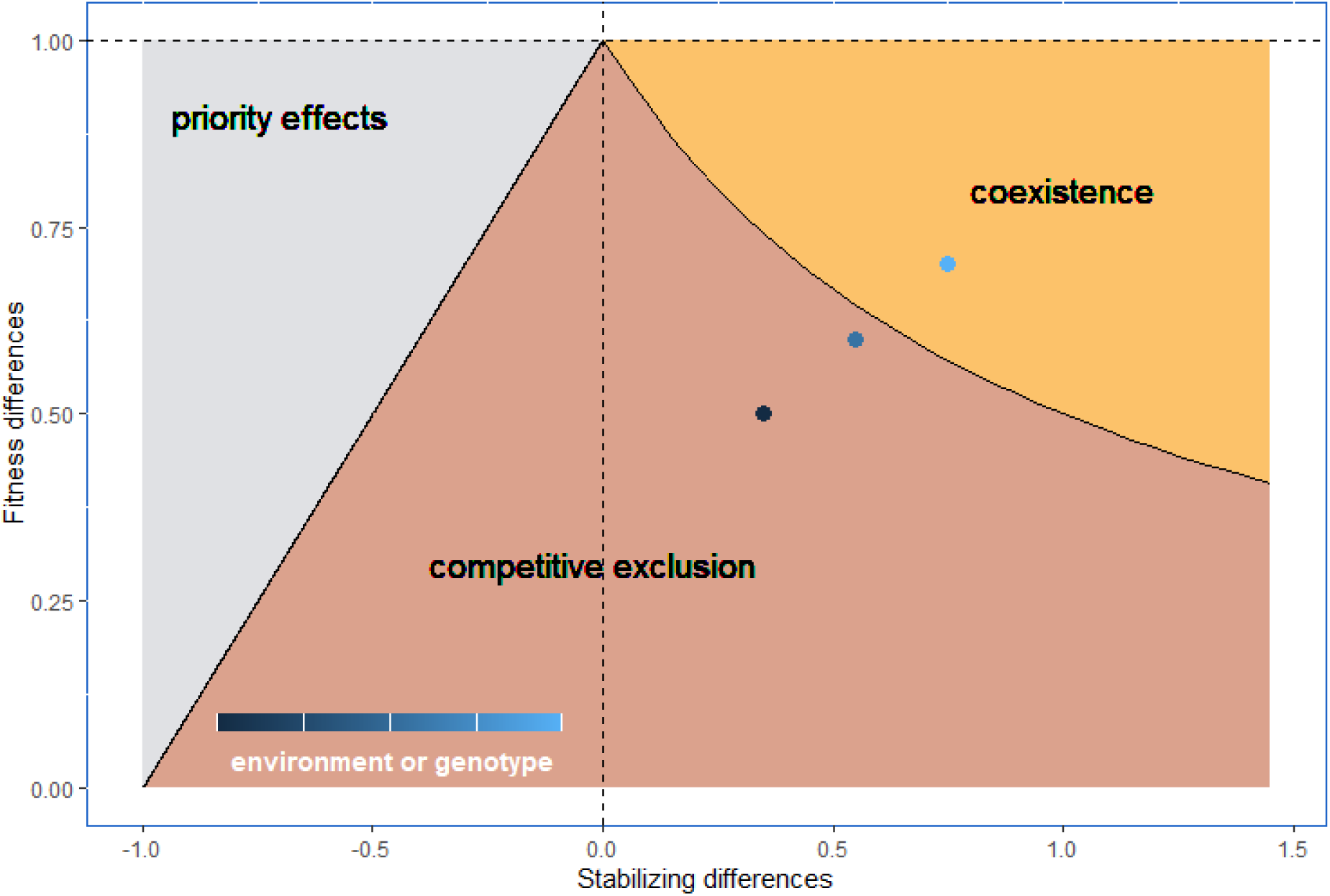
conceptual framework of modern coexistence theory adapted from (Grainger et al., 2019a) visualizing how the combination of frequency-independent effects on population growth on the y-axis, called ‘Fitness differences’, and frequency-dependent effects, called ‘Stabilizing differences’, on the x-axis determine the outcome of competition between species or cytotypes. We visualize fitness differences as the “average” fitness of the competitively inferior proportional to that of the superior. Large fitness differences lead to deterministic competitive exclusion. Negative frequency-dependency stabilizes coexistence (right) by preventing the dominance of high-proportion species/cytotypes, while positive frequency-dependency destabilizes coexistence and leads to priority effects (left). Different genotypic backgrounds or different environments can affect the outcome of competition between cytotypes (circles with different shades of blue). A more neutral outcome essentially means that species or cytotypes are similar to such an extent that fitness becomes equal (y = 1), and frequency dependency disappears (x = 0), at which point the outcome is more amenable to stochasticity.

Conversely, polyploidy provides potential avenues to improve the overall frequency independent competitiveness of the polyploid. Greater vigor and body size are anticipated to confer an immediate competitive advantage (Otto and Whitton, 2000; te Beest et al., 2012). However, evidence for polyploids as superior competitors is mixed. Polyploids have been found to be superior (Gerstein and Otto, 2011; Guo et al., 2023; Maceira et al., 1993; Sugiyama, 1998), equal (Münzbergová, 2007; Thompson et al., 2015), or inferior competitors (Anneberg et al., 2023), and even recording superior artificial polyploids but inferior natural polyploids in the same study (Castro et al., 2024). In contrast, polyploidization can also introduce frequency dependent effects on population growth. Sexual incompatibility between cytotypes, which causes MCE, has the minority cytotype encountering more incompatible gametes than the majority cytotype. MCE, therefore, results from an important positive frequency dependent effect on sexual polyploids. This depresses reproductive success for the minority cytotype, which has similar negative consequence to population growth than, for instance, effects of competition. The negative frequency dependent effects of niche differentiation are regarded as an important avenue to overcome MCE and competition with the progenitor (Fowler and Levin, 1984; Rodríguez, 1996; Thompson and Lumaret, 1992). Niche differentiation between polyploids and close relatives arises regularly by means of microgeographic differentiation (Hao et al., 2013; Mráz et al., 2012; Sonnleitner et al., 2016), stress tolerance (Anneberg et al., 2023; Bafort et al., 2023; Buggs and Pannell, 2007; Duchoslav et al., 2017; Lumaret et al., 1987; Raabová et al., 2008; Ramsey, 2007), avoiding competition for pollinators and other mutualists (Segraves and Anneberg, 2016), avoiding MCE altogether with selfing and asexuality (Chawla et al., 1997; Gustafsson, 1948; Hörandl and Hojsgaard, 2012; Miller and Venable, 2000; Van Drunen and Husband, 2018). Niche differentiation can even be caused by simple differences in the response to competition at different densities and frequencies between cytotypes (Chesson, 1994). Therefore, the effects of polyploidization on the competitive interaction of the neopolyploid with its progenitor are expected to play an important role in the establishment of polyploids, in both frequency independent and dependent effects. Stress is anticipated to be a strong mediator of these competitive interactions.

In this study, we tested the “mutual invasion” criterion in cytotype invasion experiments to understand polyploid establishment in competition with its direct diploid progenitors in replicated microcosms of the asexual *Spirodela polyrhiza* (greater duckweed). We tested the invasion of each cytotype in a population of the other cytotype for 12 weeks (>24 generations), in four neotetraploid-diploid progenitor strain pairs to accurately determine whether there is deterministic competitive exclusion of one cytotype or whether strong enough frequency dependent effects cause either stable cytotype coexistence or the certain exclusion of the invading cytotype. We compared the observed change of cytotype frequency with what is expected from differences in intrinsic growth rate alone to infer the importance of competition when testing expected polyploid establishment success. We additionally fitted population growth models to the observed invasion dynamics to estimate relative fitness and niche differences (Godwin et al., 2020). We used the classic Lotka-Volterra competition model, which incorporates intra- and interspecific competition (as developed in Chesson, 2000). This enabled us to quantify frequency dependent and independent effects, as defined within the modern coexistence framework. This theoretical framework should be especially applicable due to complete reproductive isolation. We tested invasions in control and salt-stressed conditions to infer how stress affects both cytotype’s success. Overall, we assessed to which degree competition and frequency dependent processes in mixed-ploidy populations promote polyploid success in benign and salt stressed conditions.

## Materials and Methods

### Model system and strains

Greater duckweed (*Spirodela polyrhiza*) is one of the smallest and fastest growing flowering plants. The species has a cosmopolitan distribution and thrives on freshwater surfaces that are undisturbed. Its short generation times and ease of laboratory culturing, makes it a good model system in ecology and evolution (Laird and Barks, 2018). *S. polyrhiza* also reproduces clonally, which means that the invading cytotype does not suffer from an inhibited sexual reproduction (MCE). We produced neotetraploid strains from four different diploid strains using a colchicine treatment (Hoagland’s E medium (Okunowo and Ogunkanmi, 2010) with 0.7% colchicine and 0.5% DMSO for 24h; details in Bafort et al., 2023). All neotetraploids are stable euploids. We started from four genetically different strains (9242, 9346, 9316, 0013) from the Landolt collection in Zurich, each a member of the four population genetic clusters of *S. polyrhiza* (Xu et al., 2019). All colchicine-induced neotetraploid strains are larger but show a slower intrinsic growth rate than their progenitors in benign conditions (results published in Bafort et al., 2023), similar to neopolyploids in other species (Clo and Kolář, 2021). Larger plant size may, however, help in competing for available water surface and nutrients (higher take-up rates).

### Mutual invasion microcosms

We initiated populations with ten neopolyploid fronds (i.e., individuals) of a single strain and 190 diploids of its progenitor strain to test tetraploid invasion (**tetraploid invasion** experiment). We separately initiated populations with 190 neotetraploids of a single strain with ten diploids of its progenitor strain to test diploid invasion (**diploid invasion** experiment). Each population was added to a rectangular black plastic container (surface area 14cm x 9cm, volume 1l) containing 0.2l growth medium and a transparent lid. The lid covered the container almost entirely but not completely to allow gas exchange and prevent air moisture build-up. Each week, we took a dry weight and a flow cytometry sample (see further). The remaining population was transferred to a clean black container with 0.2l fresh medium. Evaporated water was compensated by adding water (type 1) throughout the week, not exceeding the 0.2l volume to not dilute the treatment. These populations were maintained in controlled conditions (L/D 16h/8h, ±23°C).

We tested tetraploid (initial tetraploid proportion of 0.05) and diploid invasion (initial tetraploid proportion of 0.95) in four different tetraploid-diploid progenitor pairs (each given a name, 9242, 9346, 9316 and 0013, derived from the Landolt collection). We tested each pair in 2 different treatments: 1) a control treatment with Hoagland’s E medium (Okunowo and Ogunkanmi, 2010) and 2) a salt stress treatment with Hoagland’s E supplemented with 2.5 g/l NaCl. We choose a salt concentration for which earlier growth tests (in other lab conditions) found that two of our strains have a similar to higher tetraploid relative intrinsic growth rate relative to its progenitor (Bafort et al., 2023). Each combination was replicated six times. Because we expected the genotypes to remain genetically conserved, we spread the experiments in time with the 48 populations of the tetraploid invasion in the spring of 2022 and the 48 populations of the diploid invasion in the spring of 2023.

### Cytotype proportion with flow cytometry

We estimated cytotype proportion (i.e., frequency) every week. We therefore sampled 50 individuals from the population, across all life stages. To have a representative sample, we sampled in different locations of the surface after stirring the population spatial structure. These 50 individuals implied the removal of a small fraction of the total population size that quickly grew to well over 1000 individuals in all populations. Each sample of 50 individuals was processed as two mixed-ploidy flow cytometry samples of 25 individuals in the tetraploid invasion, whose counts were then added together. By the start of the diploid invasion experiment, we found that all 50 individuals could be processed as one mixed-ploidy flow cytometry sample in the diploid invasion and still get clean flow cytometry results. The individuals were chopped together in a petri dish with a razor blade for 1 min in an adapted Galbraith buffer (supplementary materials, Appendix 1) and filtered (40µm) to extract nuclei. The extracted nuclei were preserved on ice until stained with DAPI (4µg/ml). We followed the one-step protocol of nuclei extraction and flow cytometry of Dolezel et al. (2007). Stained samples were processed by the flow cytometer (Attune NxT Acoustic Focusing Cytometer) and ploidy was determined by external controls of known diploid and tetraploid *S. polyrhiza* samples. We extracted the number of measured diploid nuclei by gating a fixed size range on the DAPI-detecting violet axis, centered on the peak that was found around the value where diploid nuclei were expected from the external control. Tetraploid nuclei were extracted from a gate with lower and upper bound twice the value of the respectively lower and upper bound of the diploid gate. Flow cytometric results were analyzed using FlowJo (BD Life Sciences, 2023).

The proportion of tetraploid nuclei provides a good approximation but is not exactly equal to the proportion of tetraploid individuals in the sample. This has three main reasons. 1) A small proportion of nuclei have double the ploidy of its individual because of endopolyploidy or of cells in G2 of the cell cycle, i.e., after DNA replication but before cell division. 2) Neotetraploids in our experimental system are on average bigger and have bigger cells than plants from their progenitor diploid population. A neotetraploid individual may therefore provide a different number of nuclei compared to a diploid. Though, it is unclear whether an average neotetraploid individual has more, less, or the same number of cells than an average diploid. 3) Stained or autofluorescent debris particles that are not nuclei are counted within diploid and tetraploid gates. To correct for all three effects, we constructed a calibration data set by measuring a number of samples with known tetraploid proportion for each neotetraploid-diploid progenitor pair using the same procedure as for the experimental samples. We measured one sample of 48 individuals for each neotetraploid-diploid pair that had either 2, 4, 6, 8, 10, 12, 16, 20, 24, 28, 32, 36, 38, 40, 42, 44 or 46 diploids. Calibration curves can be found in Appendix 2. The measured individuals were grown in control conditions (Hoagland medium without salt) and low densities, but we assume that the calibration needed is similar for duckweed grown in other conditions. These samples containing known diploid and tetraploid individuals from the four different strains also functioned as an external control to determine diploid nuclei peaks in the experimental data. Nuclei fluorescence on the violet axis were stable enough to reliably distinguish diploid from tetraploid peaks across all 12 weeks.

### Dry weight

Each week, we measured dry weight to observe absolute population growth. We collected in each replicate all duckweed in a circle of radius 1.75cm (area: 9.62 cm²) from the 14cm x 9cm of the box (area: 126 cm², 7.6% of the population removed for measurement each week), making sure to have all layers of duckweed growing on top of each other in that area. We stored the samples in small envelopes at -10°C for weighing later. All samples were dried in those envelopes for 5 days in an 80°C oven with ventilation. Pilot experiments showed that this duration was sufficient to evaporate most moisture from the sample. Each sample was then weighed on a microbalance within 1 minute of leaving the oven to prevent much air moisture take-up of the envelopes or dry plant tissue. Further details on the model description and results are provided in the supplementary materials (Appendix 3).

### Population weight to size conversion

To estimate the weight per individual, we measured dry weight of 96 samples with a known number of duckweed individuals from both ploidies in all four strains. We tested three replicates for 100, 250, 500, and 1000 individuals using the same dry weight procedure as for the experimental population samples. We used pre-weighed aluminum envelopes to be able to calculate plant dry weight exactly. Further details on the model description and results are provided in the supplementary materials (Appendix 4).

### Statistical analyses

In the experiment, we did not measure the tetraploid individual proportion (*p*4*n*_*ind*_) but the tetraploid nuclei proportion (*p*4*n*_*nuclei*_). However, we assume that growth medium treatment, strain, and time affect the individual proportion directly, not the nuclei. Therefore, we first estimated the experimental individual proportion (*p*4*n*_*ind*_) from the measured tetraploid nuclei proportion (*p*4*n*_*nuclei*_) using latent-variable imputation. We modelled the logit-transformed tetraploid nuclei proportion as a response with normal error distribution (1) that is affected by the tetraploid individual proportion according to a strain-specific intercept and slope (2). By inferring strain-specific intercepts (𝛼_[𝑠𝑡𝑟𝑎*in*]_) and slopes (𝛽_[𝑠𝑡𝑟𝑎*in*]_) from the calibration data, we can impute the unknown tetraploid individual proportion in the experimental populations from their measured tetraploid nuclei proportion. We logit-transformed proportions (i.e., log-odds), a common practice when inferring effects on a proportion. This method estimates a posterior distribution of tetraploid individual proportion for each measured tetraploid (nuclei) proportion in the experimental dataset.

We infer strain and environment effects based on 50 datasets sampled from the posterior distribution of tetraploid individual proportion for each experimental data point. We modelled logit-transformed tetraploid individual proportion as a response variable with a normal error distribution (3). We modelled the proportion of tetraploid individuals as the response of linear effects of strain, growth medium and their interaction (𝛾_[𝑠𝑡𝑟𝑎*in* 𝑋 𝑠𝑎*l*𝑡]_) on the change of proportion across time (week). We modelled a fixed intercept at proportion 0.05 (𝐼*n*𝑡*e*𝑟*cep*𝑡 = logit (0.05) = -2.94) for the tetraploid invasion and proportion 0.95 (𝐼*n*𝑡*e*𝑟*cep*𝑡 = logit (0.95) = 2.94) for the diploid invasion to reflect the starting tetraploid proportions in the experiment (4). We also model a variable slope in time per replicate population (pop) to account for population-specific differences in tetraploid frequency change (5). Both models can be formulated as follows:

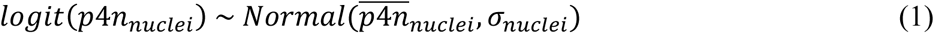

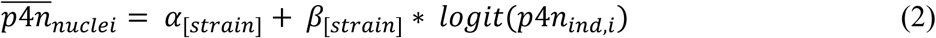

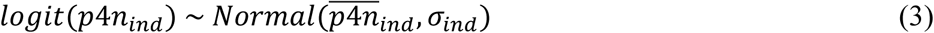

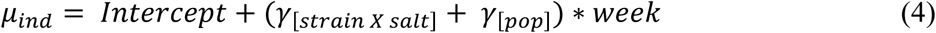

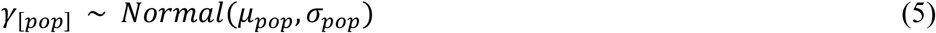

To evaluate the effect of fixing the intercept, we provide results of the same model except for estimating the effects of strain, salt, and population on the intercept as well (supplementary materials Appendix 2, 5).

There are limits to the accuracy of estimating the tetraploid proportion with this method. The logit transformation causes inaccuracies of the flow cytometry (in experiment and calibration) to have a proportionally bigger effect on very low and very high proportions. The calibration model deals with this uncertainty by avoiding overcorrection and, therefore, overestimating very low proportions and underestimating very high proportions. This makes it hard to detect the complete extinction of tetraploids or diploids.

We compared the observed tetraploid proportion from our experiment with tetraploid proportion trajectories that are expected from differences in intrinsic growth (i.e., exponential growth). Details on how we measured intrinsic growth and the underlying model assumed can be found in supplementary materials (Appendix 6)

We, then, estimate tetraploid and diploid population sizes from combining the posterior distributions of experimental tetraploid proportions, experimental dry weight (Appendix 2), and individual dry weight (Appendix 3). Because of error propagation over the three datasets, population size estimates have a large uncertainty envelope. This showed to be especially true when estimating the overall low tetraploid proportion in the tetraploid invasion experiment. We therefore restricted these analyses to the quantification of population size in the diploid invasion experiment. Each combination of 50 posterior draws of these three model posteriors are used to calculate expected total population size (𝑁_𝑡𝑜𝑡𝑎*l*_) for each experimental data point (i) as follows:

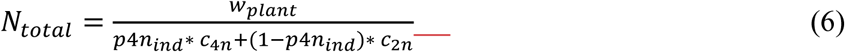

With 𝑤_*pl*𝑎*n*𝑡_the estimated total plant dry weight for each data point, *p*4*n*_*ind*_the estimated tetraploid proportion of each data point and *c*_*i*_ the (strain-specific) individual weight for cytotype i. The denominator is the average individual weight weighted according to the cytotype proportion. Cytotype-specific population sizes are easily calculated: 𝑁_4*n*_ = *p*4*n*_*ind*_ ∗ *n*_𝑡𝑜𝑡𝑎*l*_ and 𝑁_2*n*_ = (1 − *p*4*n*_*ind*_) ∗ 𝑁_𝑡𝑜𝑡𝑎*l*_.

These series of estimated population sizes were subsequently fitted to recursive differential equations of Lotka-Volterra for population growth of competing species/cytotypes. In doing so, we estimated interaction coefficients (𝛼) from the population size time-series for both cytotypes:

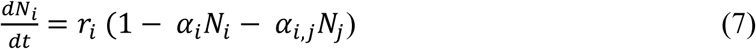

With 𝑁_*i*_the population size of cytotype i, 𝑁_𝑗_ the population size of its competitor cytotype, 𝑟_*i*_the per-week intrinsic growth rate of cytotype i as estimated from our monoculture growth tests (cfr. 6), 𝛼_*i*_ the competition coefficient of cytotype i on its own population growth (intracytotype competition coefficient) and 𝛼_*i*,𝑗_the competition coefficient of the competitor cytotype on cytotype i’s population growth. We use initial population sizes of 10 diploids and 190 tetraploids in the diploid invasion experiment. A more detailed model description and estimated coefficients can be found in supplementary materials (Appendix 7). The four competition coefficient (intracytotype: 𝛼_2*n*_, 𝛼_4*n*_; intercytotype: 𝛼_2*n*,4*n*_, 𝛼_4*n*,2*n*_) are then combined to quantify the frequency independent component affecting competition outcomes (8) and the frequency dependent component affection competition outcomes (9), as derived by modern coexistence theory (Chesson, 2018):

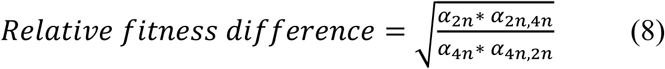

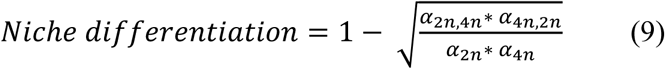

All analyses were performed in R (R core team, 2021) using the brms library (Bürkner, 2018) for Bayesian statistics that is based on Stan (Stan Development Team, 2019) and the tidyverse ecosystem (Wickham et al., 2019). Bayesian inference was performed using Hamiltonian Monte Carlo (HMC) models that implemented four chains with each 4,000 iterations from which 2,000 were considered burn-in. The HMC model to fit the recursive population models utilized the Dormand-Prince ODE solver in Stan with additional absolute and relative tolerance of 0.001 and maximum number of steps between timepoints of 100. Priors were used that are weakly regularizing by choosing prior distributions that are significantly wider than the parameter values that would be reasonable to expect for each model. We evaluated the performance of every fitted model based on standardized procedures by checking mixing and stationarity in the trace plots and by checking the effective sample size and 𝑅^ statistic for each parameter (McElreath, 2020). Models, formulae, and detailed description of the procedure and priors can be found in the electronic supplementary material.

## Result

Average tetraploid proportion decreased in all strain-treatment combinations (Fig. 2, marginal slopes in Fig. 3) in both the tetraploid and diploid invasion experiment. In the tetraploid invasion experiment, tetraploid proportion showed a remarkably similar slight decline for all strain-treatment combinations (dashed violins, Fig. 3). The diploid invasion experiment shows a faster and consistent tetraploid decline over all strain-environment combinations. The salt treatment had a different effect on tetraploid proportion decline in different strains (Fig. 2). Moreover, we found substantial variability in proportion trajectories within each genotype-treatment combination (Fig. S8).

**Fig. 2:**
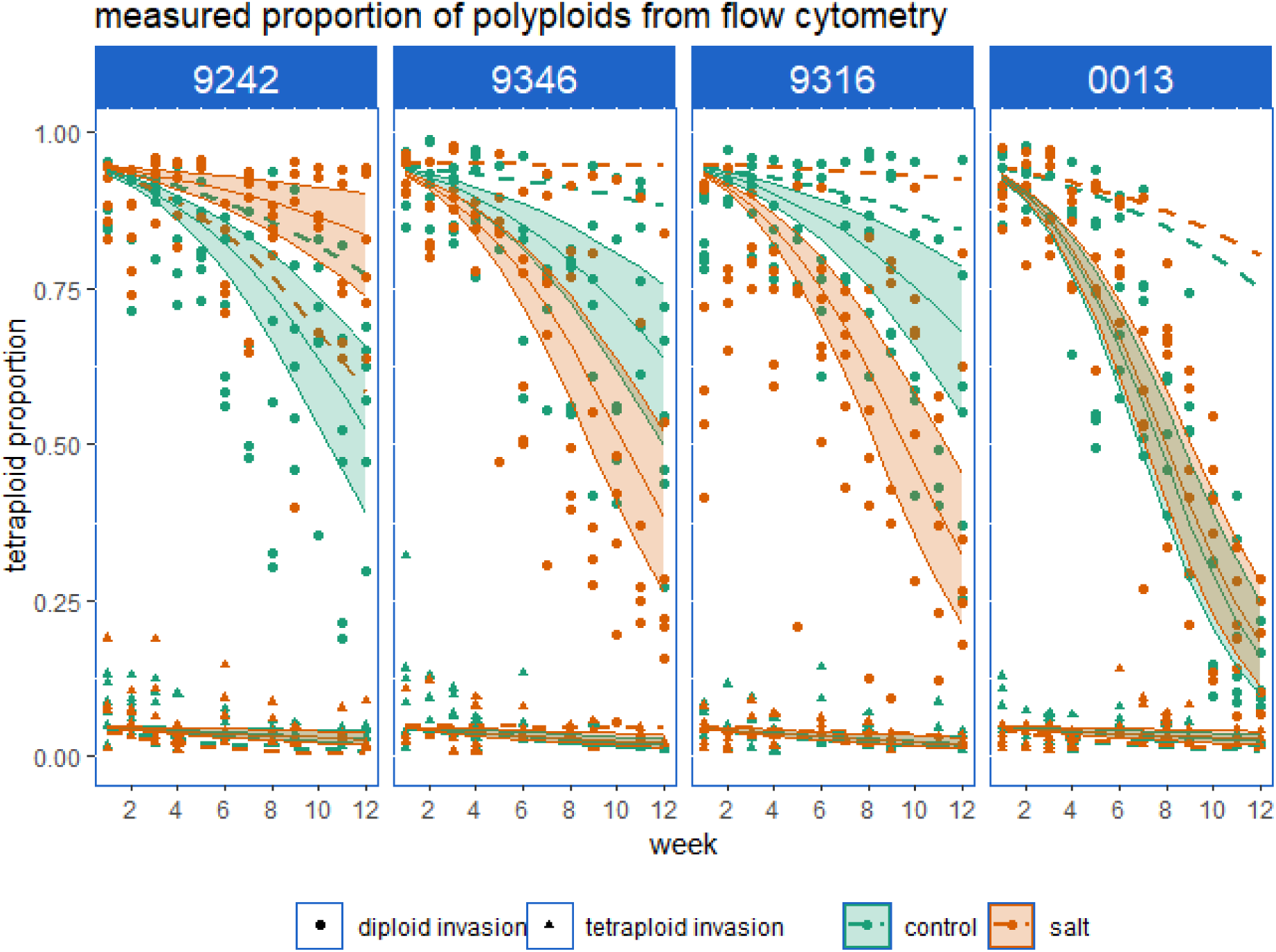
Tetraploid proportion during tetraploid invasion (triangles) and diploid invasion (circles) experiment for different strains in a control (green) or salt stress (orange) environment. Lines and ribbons represent the mean and [0.09, 0.91] likelihood interval, respectively, of the posterior expected prediction. Dashed lines indicate expectations from differences in intrinsic growth rate estimated in separate growth tests.

**Fig. 3:**
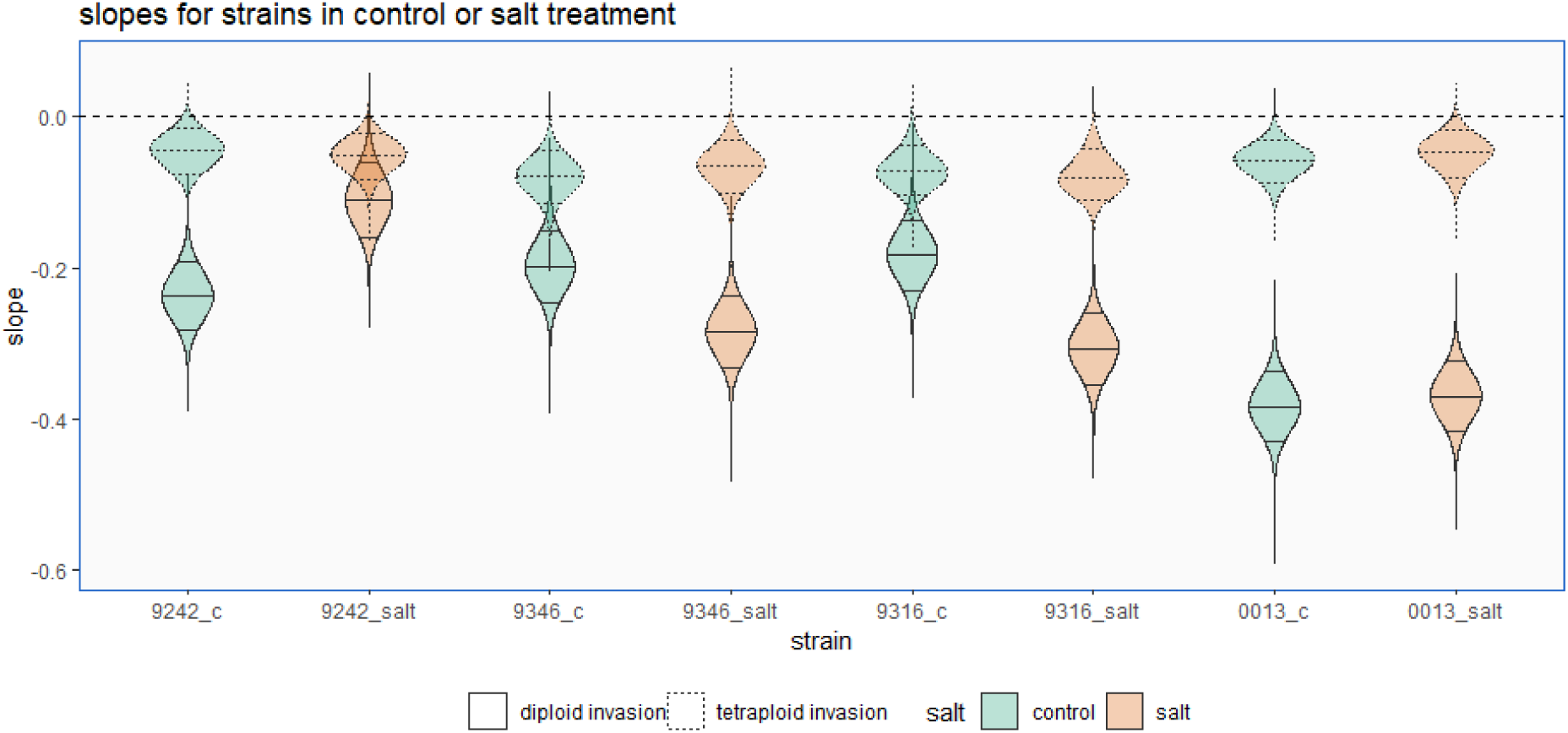
Tetraploid proportion decline from posterior distributions of marginal slopes of all strains (9242, 9346, 9316, 0013), in control (green) or salt (orange) medium for the tetraploid invasion (dotted violins) and diploid invasion (full violin) experiments. We indicate the median, 9th and 91st percentile in each violin.

The comparison of the observed changes in tetraploid proportion to the expected change assuming exponential growth indicated the prevalence of substantial competition effects (Fig. 2, dashed lines). Hence, tetraploid invasion success over the progenitor cannot be accurately predicted from differences in intrinsic growth rate alone. Except for strain 9242 in salt, all diploid invasions proceeded faster than expected from differences in intrinsic growth rate alone, which suggests that tetraploids are more sensitive to competition relative to diploids, or that negative frequency dependent growth disproportionately favor the invading diploids.

Population dynamics as derived from the recursive differential equations for population growth of both cytotypes demonstrate the pertinent invasion ability of diploids in all strain-treatment combinations, or conversely a stagnation to decline of tetraploid population growth (Fig. 4). Given the most likely parameter estimates, stabilizing negative frequency dependent growth, i.e., niche differentiation, prevail across the experiment (Fig. 5). The trajectories for strain 9242 and 9346 are even estimated to have enough negative frequency dependent growth to overcome the fitness differences with an expected stable coexistence. This contrasts to the tetraploid invasion experiment where tetraploids of both strains were unable to grow in proportion during the experimental timeframe (Fig. 2).

**Fig. 4:**
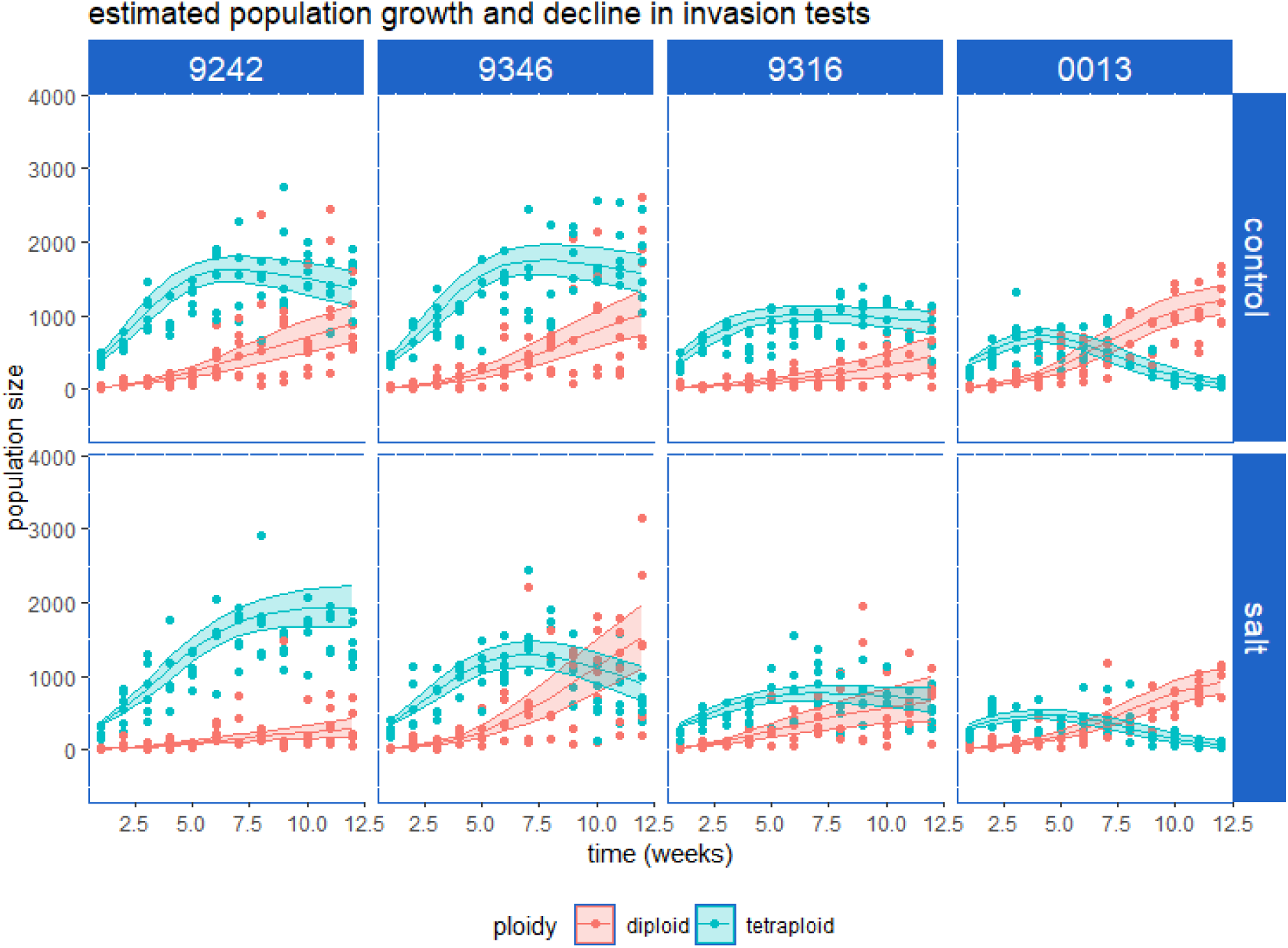
Estimated population size (in number of individuals) of diploids (red) and tetraploids (indigo) averaged over all replicates for each strain-stress treatment combination in the diploid invasion experiment. Lines and ribbons represent the mean and [0.09, 0.91] likelihood interval, respectively, of the posterior expected prediction.

**Fig. 5:**
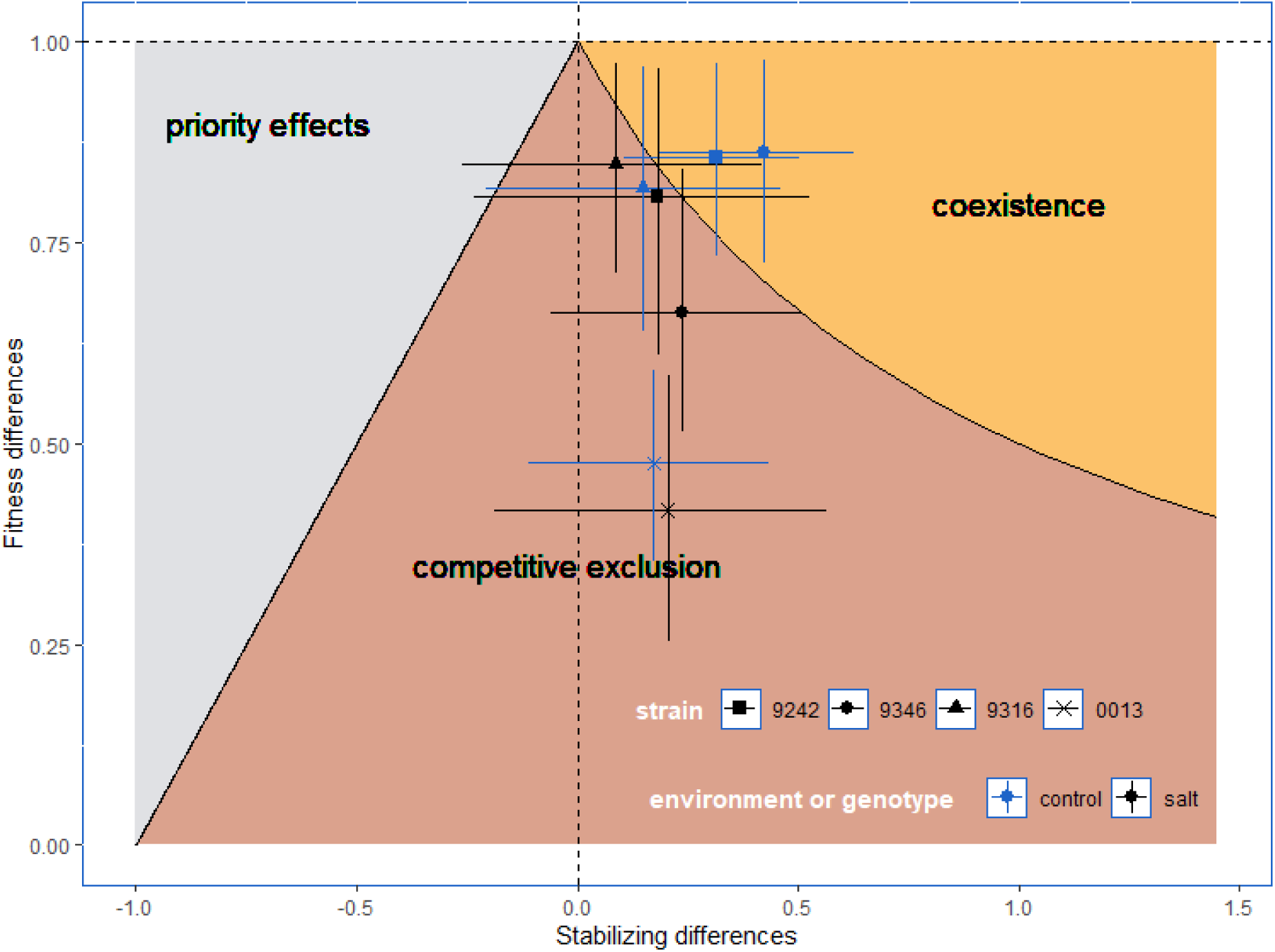
Estimated stabilizing differences (niche differentiation) and fitness differences from estimated parameters of the recursive population models, averaged over all replicates for each strain-treatment combination. We also plot the [0.09, 0.91] likelihood intervals to show the sizeable uncertainty in these parameters. Figure adapted from Grainger et al. (2019a).

## Discussion

Mutual invasion experiments showed that selected strains of neotetraploid *Spirodela polyrhiza* were not able to invade their diploid progenitor populations but that the diploid progenitor was clearly able to invade neotetraploid populations in all strain-salt combinations. However, the similar, and slow, decline of tetraploid frequency in tetraploid invasion experiment was likely caused by an underestimation of decline due to the inaccurate estimation of lower proportions. Despite the underestimation of tetraploid proportion decline, we still confidently conclude a decline by the way the change was consistent enough for the model to estimate almost exclusively negative posterior slopes for all strain-treatment combinations (dashed violins, Fig. 3). Overall, a consistent lack of tetraploid success was clear in all strain-treatment combinations across 12 weeks. Invasion speed, estimated by the rate of tetraploid proportion increase or decline, differed clearly between different strains with no consistent relative benefit or harm of salt stress on tetraploid success across the studied strains. Coexistence of both cytotypes may, however, be possible at longer time frames based on the estimated negative frequency dependency in some diploid-polyploid strains.

Polyploid *S. polyrhiza* populations failed to invade their progenitor population and were successfully invaded by their progenitor. This is in line with the overall lower intrinsic growth rates of neotetraploids relative to diploid progenitors in these (Bafort et al., 2023) and other neopolyploid *S. polyrhiza* strains (Anneberg et al., 2023). The failed invasion of neopolyploids indicates that all neopolyploid strains faced a deterministic competitive exclusion in both control and salt-stressed conditions (brown zone in Fig. 1, 5). We assessed invasion dynamics in a direct way, which is different to an indirect inference of competition outcomes from measuring short-term fitness differences in different competitor densities or by measuring the short-term invasion rate of an invader into an established population (Godwin et al., 2020; Grainger et al., 2019b). However, a non-simplified assessment of the invasion dynamics avoids possible errors, for instance from carry-over effects on growth rate in the initial period of invasion when measured.

Despite being quite certain that tetraploid proportions declined, we found that they declined at a remarkably similar (low) rate in the tetraploid invasion experiment in all strain-treatment combinations. This exposes the weakness in our method of tetraploid proportion estimation from mixed-cytotype flow cytometry. Accurate estimation of low tetraploid proportions is hindered because of tetraploid nuclei produced from diploid duckweed (due to the cell cycle or endopolyploidy) and autofluorescent debris. These particles that pose as nuclei from tetraploids outnumber the “real” tetraploid nuclei from tetraploid individuals in the flow cytometer at low tetraploid proportion. Calibration helps to correct for these biases, but the additional uncertainty results in the underestimation of tetraploid proportion decline at low proportions. Therefore, we only draw further conclusions from the diploid invasion experiment.

The diploid invasion trajectories that we observed indicated a faster diploid invasion than what we would expect from differences in intrinsic growth rate alone. Competition showed to even facilitate diploid invasions. Because we can only draw reliable conclusions from the diploid invasion experiment, we cannot discern whether diploid invasions were enabled because of a (frequency independent) higher sensitivity of the tetraploid to competition or that negative frequency dependent growth benefitted the lower frequency diploids. In any case, intrinsic growth rate estimations are thus not reliable indicators for polyploid success.

We also estimated frequency-dependency and predicted the outcome of competition by means of the modern coexistence theory (Chesson, 2000; Godwin et al., 2020). The estimated parameter combinations confirmed the retrieved competitive exclusion of the tetraploid in six of eight strain-treatment combinations (Fig. 5) and showed negative frequency dependent growth in all eight strain-treatment combinations. This niche differentiation was estimated to be sufficiently strong to expect stabile coexistence between neopolyploids and progenitors in two strains in the control medium. The expectation of stable coexistence contradicts the lack of a successful tetraploid invasion in the tetraploid invasion experiment, which would be expected for stable coexistence. Wide error bars, however, indicate rough estimations because they were made after several data models, each propagating their inaccuracies. The time-series were likely too short to have an accurate estimation of the competition coefficients used to calculate the frequency independent and dependent parameters. While we show what is possible in interpreting polyploid establishment in the light of the modern coexistence framework, we also show that different methods lead us to draw different conclusions. The framework of modern coexistence theory, while capturing an important aspect of population success, must be applied to observed populations with great care for the underlying assumptions.

The rate of tetraploid proportion decline varied notably among strains and treatments. Neotetraploid species commonly show a decreased performance (Clo and Kolář, 2021) but with sizeable differences between genotypes (strains, observed in, e.g., *Spirodela polyrhiza* in Anneberg et al., 2023; Bafort et al., 2023). In general, an observed interaction of WGD and genetic background may stem from an interaction with the ancestral genotype or from stochastic processes during WGD that affect the polyploid genotype. Indeed, the ancestral genotype may affect, or be affected by, changes related to WGD. For instance, The genotype may alter the magnitude of cell and body size increase (as seen in Bafort et al., 2023) or by experiencing different gene dosage effects (Adams and Wendel, 2005; Doyle and Coate, 2019). However, stochastic (epi)genetic or genomic alterations that happen at or close after WGD may also underlie differences in performance (called ‘genome shock’ for allopolyploids, Shimizu, 2022; but also a factor in autopolyploids Liu et al., 2017; Parisod et al., 2010; Tayalé and Parisod, 2013). We do not know the extent of (epi)-genetic and genomic alteration in our neotetraploid and diploid progenitors *S. polyrhiza* strains but presume few genomic deletions based on cytometric data. Because each tetraploid strain is the result of a single independent WGD from a different strain, we cannot separate the effects of ancestral genotype from the effects of WGD-related (epi)genomic stochasticity.

Our experimental invasion dynamics also showed strong variation within strain and treatments (Fig. S8). This demographic stochasticity clearly had a lasting effect on the population size trajectory due to the compounding effects of differences in earlier generations on later generations. A lack of strong frequency-dependent and -independent effects means that stochasticity can have a relevant effect on the frequency of competitors (Adler et al., 2007; Grainger et al., 2019a). It may explain the accidental invasion of non-adaptive diploid yeast *Saccharomyces cerevisiae* in haploid experimental evolution populations (Gerstein and Otto, 2011). The impact of demographic stochasticity is widely recognized for the establishment of species in a community (Hubbell, 2001) or genetic variants in a population (Kimura, 1983; Ohta and Gillespie, 1996), and should not be underestimated when thinking about polyploid establishment.

In conclusion, we found that neotetraploid strains of greater duckweed *Spirodela polyrhiza* were unable to invade their progenitor strain and were invaded by their progenitor strain. The tetraploid proportion decreased at a different rate across different strains (genotypic backgrounds) and environments (control compared to salt stress) but not sufficiently to overcome inherent neotetraploid disadvantages. Rough estimations of recursive population models on our experimental time-series indicated that there is likely negative frequency dependency between neotetraploids and their progenitor present in all strain-treatment combinations.

## Acknowledgements

We thank Annelore Natran, Michiel Kesteleyn, Noa Bauwens and Liam Aelvoet who helped with the competition experiment transfers and flow cytometry procedures. We also thank Walter Lämmler and the Landolt collection for providing us with biological material, the staff of the VIB flow Core for their assistance with flow cytometry, Quinten Bafort and Tian Wu for providing the neotetraploid and progenitor strains and expertise with the model system, and Maxime Dahirel and Arthur Zwaenepoel for advice on the statistical modelling. FM was supported by Fonds Wetenschappelijk Onderzoek (FWO, 1238124N). This work was also supported by the European Research Council under the European Union’s Horizon 2020 Research and Innovation program (No. 833522) and from Ghent University (Methusalem funding, BOF.MET.2021.0005.01, to YVdP).

## Author contributions

FM, YVdP and DB designed the experiment, FM collected data and analyzed the data, FM, YVdP and DB interpreted the data and wrote this manuscript.

## Data availability

Data and R code to analyze the data and generate figures are available at https://github.com/fremorti/Spirodela-invasion-experiment-to-test-establishment

## References

1. Adams, K.L., Wendel, J.F., 2005. Polyploidy and genome evolution in plants. Curr. Opin. Plant Biol. 8, 135–141. 10.1016/j.pbi.2005.01.001

2. Adler, P.B., HilleRisLambers, J., Levine, J.M., 2007. A niche for neutrality. Ecol. Lett. 10, 95–104. 10.1111/j.1461-0248.2006.00996.x

3. Anneberg, T.J., O’Neill, E.M., Ashman, T.L., Turcotte, M.M., 2023. Polyploidy impacts population growth and competition with diploids: multigenerational experiments reveal key life-history trade-offs. New Phytol. 1294–1304. 10.1111/nph.18794

4. Bafort, Q., Wu, T., Natran, A., De Clerck, O., Van De Peer, Y., 2023. The immediate effects of polyploidization of Spirodela polyrhiza change in a strain-specific way along environmental gradients. Evol. Lett. 7, 37–47. 10.1093/evlett/qrac003

5. Barabás, G., D’Andrea, R., Stump, S.M., 2018. Chesson’s coexistence theory. Ecol. Monogr. 88, 277–303. 10.1002/ecm.1302

6. BD Life Sciences, 2023. FlowJo Software.

7. Bomblies, K., 2020. When everything changes at once: Finding a new normal after genome duplication: Evolutionary response to polyploidy. Proc. R. Soc. B Biol. Sci. 287. 10.1098/rspb.2020.2154

8. Buggs, R.J.A., Pannell, J.R., 2007. ECOLOGICAL DIFFERENTIATION AND DIPLOID SUPERIORITY ACROSS A MOVING PLOIDY CONTACT ZONE. Evolution 61, 125–140. 10.1111/j.1558-5646.2007.00010.x

9. Bürkner, P.-C., 2018. Advanced Bayesian Multilevel Modeling with the R Package brms. R J. 10, 395. 10.32614/RJ-2018-017

10. Castro, M., Celeste Dias, M., Loureiro, J., Husband, B.C., Castro, S., 2024. Competitive ability, neopolyploid establishment and current distribution of a diploid–tetraploid plant complex. Oikos 2024, e09949. 10.1111/oik.09949

11. Chase, J.M., Leibold, M.A., 2003. Ecological Niches. University of Chicago Press. 10.7208/chicago/9780226101811.001.0001

12. Chawla, B., Bernatzky, R., Liang, W., Marcotrigiano, M., 1997. Breakdown of self-incompatibility in tetraploid Lycopersicon peruvianum: Inheritance and expression of S-related proteins. Theor. Appl. Genet. 95, 992–996. 10.1007/s001220050652

13. Chesson, P., 2018. Updates on mechanisms of maintenance of species diversity. J. Ecol. 106, 1773–1794. 10.1111/1365-2745.13035

14. Chesson, P., 2000. Mechanisms of Maintenance of Species Diversity. Annu. Rev. Ecol. Syst. 31, 343–366. 10.1146/annurev.ecolsys.31.1.343

15. Chesson, P., 1994. Multispecies Competition in Variable Environments. Theor. Popul. Biol. 45, 227–276. 10.1006/tpbi.1994.1013

16. Clo, J., Kolář, F., 2021. Short- and long-term consequences of genome doubling: a meta-analysis. Am. J. Bot. 108, 2315–2322. 10.1002/ajb2.1759

17. Comai, L., 2005. The advantages and disadvantages of being polyploid. Nat. Rev. Genet. 6, 836–846. 10.1038/nrg1711

18. De Meester, L., Vanoverbeke, J., Kilsdonk, L.J., Urban, M.C., 2016. Evolving Perspectives on Monopolization and Priority Effects. Trends Ecol. Evol. 31, 136–146. 10.1016/j.tree.2015.12.009

19. Doležel, J., Greilhuber, J., Suda, J., 2007. Estimation of nuclear DNA content in plants using flow cytometry. Nat. Protoc. 2, 2233–2244. 10.1038/nprot.2007.310

20. Doyle, J.J., Coate, J.E., 2019. Polyploidy, the nucleotype, and novelty: The impact of genome doubling on the biology of the cell. Int. J. Plant Sci. 180, 1–52. 10.1086/700636

21. Duchoslav, M., Fialová, M., Jandová, M., 2017. The ecological performance of tetra-, penta- and hexaploid geophyte Allium oleraceum in reciprocal transplant experiment may explain the occurrence of multiple-cytotype populations. J. Plant Ecol. 10, 569–580. 10.1093/jpe/rtw039

22. Felber, F., 1991. Establishment of a tetraploid cytotype in a diploid population: Effect of relative fitness of the cytotypes. J. Evol. Biol. 4, 195–207. 10.1046/j.1420-9101.1991.4020195.x

23. Fowler, N.L., Levin, D.A., 1984. Ecological constraints on the establishment of a novel polyploid in competition with its diploid progenitor. Am. Nat. 124, 703–711. 10.1086/284307

24. Fukami, T., 2015. Historical Contingency in Community Assembly: Integrating Niches, Species Pools, and Priority Effects. Annu. Rev. Ecol. Evol. Syst. 46, 1–23. 10.1146/annurev-ecolsys-110411-160340

25. Gause, G.F., 1934. The struggle for existence. The Williams & Wilkins company, Baltimore, 10.5962/bhl.title.4489

26. Gaynor, M.L., Kortessis, N., Soltis, D.E., Soltis, P.S., Ponciano, J.M., 2024. Dynamics of mixed-ploidy populations under demographic and environmental stochasticities [WWW Document]. 10.1086/734411. https://doi.org/10.1086/734411

27. Gerstein, A.C., Otto, S.P., 2011. Cryptic Fitness Advantage: Diploids Invade Haploid Populations Despite Lacking Any Apparent Advantage as Measured by Standard Fitness Assays. PLoS ONE 6, e26599. 10.1371/journal.pone.0026599

28. Godwin, C.M., Chang, F.-H., Cardinale, B.J., 2020. An empiricist’s guide to modern coexistence theory for competitive communities. Oikos 129, 1109–1127. 10.1111/oik.06957

29. Grainger, T.N., Letten, A.D., Gilbert, B., Fukami, T., 2019a. Applying modern coexistence theory to priority effects. Proc. Natl. Acad. Sci. 116, 6205–6210. 10.1073/pnas.1803122116

30. Grainger, T.N., Levine, J.M., Gilbert, B., 2019b. The Invasion Criterion: A Common Currency for Ecological Research. Trends Ecol. Evol. 34, 925–935. 10.1016/j.tree.2019.05.007

31. Guignard, S., Nichols, R.A., Knell, R.J., Macdonald, A., Romila, C., Trimmer, M., Leitch, I.J., Leitch, A.R., 2016. Genome size and ploidy influence angiosperm species ’ biomass under nitrogen and phosphorus limitation. New Phytol. 210, 1195–1206.

32. Guo, W., Wei, N., Hao, G.Y., Yang, S.J., Zhu, Z.Y., Yang, Y.P., Duan, Y.W., 2023. Does competitive asymmetry confer polyploid advantage under changing environments? J. Ecol. 1–13. 10.1111/1365-2745.14100

33. Gustafsson, Å., 1948. POLYPLOIDY, LIFE-FORM AND VEGETATIVE REPRODUCTION. Hereditas 34, 1–22. 10.1111/j.1601-5223.1948.tb02824.x

34. Hao, G.Y., Lucero, M.E., Sanderson, S.C., Zacharias, E.H., Holbrook, N.M., 2013. Polyploidy enhances the occupation of heterogeneous environments through hydraulic related trade-offs in Atriplex canescens (Chenopodiaceae). New Phytol. 197, 970–978. 10.1111/nph.12051

35. Hörandl, E., Hojsgaard, D., 2012. The evolution of apomixis in angiosperms: A reappraisal. Plant Biosyst. 146, 681–693. 10.1080/11263504.2012.716795

36. Hubbell, S.P., 2001. The Unified Neutral Theory of Biodiversity and Biogeography. Princeton university Press.

37. Husband, B.C., 2000. Constraints on polyploid evolution: A test of the minority cytotype exclusion principle. Proc. R. Soc. B Biol. Sci. 267, 217–223. 10.1098/rspb.2000.0990

38. Husband, B.C., Sabara, H.A., 2004. Reproductive isolation between autotetraploids and their diploid progenitors in fireweed, Chamerion angustifolium (Onagraceae). New Phytol. 161, 703–713. 10.1046/j.1469-8137.2004.00998.x

39. Irwin, D., Schluter, D., 2022. Hybridization and the Coexistence of Species. Am. Nat. 200, E93–E109. 10.1086/720365

40. Kauai, F., Mortier, F., Milosavljevic, S., Van de Peer, Y., Bonte, D., 2023. Neutral processes underlying the macro eco-evolutionary dynamics of mixed-ploidy systems. Proc. R. Soc. B 290. 10.1098/rspb.2022.2456

41. Kimura, M., 1983. The Neutral Theory of Molecular Evolution. Cambridge University Press.

42. Laird, R.A., Barks, P.M., 2018. Skimming the surface: duckweed as a model system in ecology and evolution. Am. J. Bot. 105, 1962–1966. 10.1002/ajb2.1194

43. Levin, D.A., 1975. MINORITY CYTOTYPE EXCLUSION IN LOCAL PLANT POPULATIONS. TAXON 24, 35–43. 10.2307/1218997

44. Liu, S., Yang, Y., Wei, F., Duan, J., Braynen, J., Tian, B., Cao, G., Shi, G., Yuan, J., 2017. Autopolyploidy leads to rapid genomic changes in Arabidopsis thaliana. Theory Biosci. 136, 199–206. 10.1007/s12064-017-0252-3

45. Lumaret, R., Guillerm, J.-L., Delay, J., Ait Lhaj Loutfi, A., Izco, J., Jay, M., 1987. Polyploidy and habitat differentiation in Dactylis glomerata L. from Galicia (Spain). Oecologia 73, 436–446. 10.1007/BF00385262

46. Maceira, N.O., Jacquard, P., Lumaret, R., 1993. Competition between diploid and derivative autotetraploid Dactylis glomerata L. from Galicia. Implications for the establishment of novel polyploid populations. New Phytol. 124, 321–328. 10.1111/j.1469-8137.1993.tb03822.x

47. McElreath, R., 2020. Statistical rethinking: A Bayesian course with examples in R and Stan. Chapman and Hall/CRC press.

48. Miller, J.S., Venable, D.L., 2000. Polyploidy and the evolution of gender dimorphism in plants. Science 289, 2335–2338. 10.1126/science.289.5488.2335

49. Mortier, F., Bafort, Q., Milosavljevic, S., Kauai, F., Prost Boxoen, L., Van de Peer, Y., Bonte, D., 2024. Understanding polyploid establishment: temporary persistence or stable coexistence? Oikos e09929. 10.1111/oik.09929

50. Mráz, P., Španiel, S., Keller, A., Bowmann, G., Farkas, A., Šingliarová, B., Rohr, R.P., Broennimann, O., Müller-Schärer, H., 2012. Anthropogenic disturbance as a driver of microspatial and microhabitat segregation of cytotypes of Centaurea stoebe and cytotype interactions in secondary contact zones. Ann. Bot. 110, 615–627. 10.1093/aob/mcs120

51. Münzbergová, Z., 2007. No effect of ploidy level in plant response to competition in a common garden experiment. Biol. J. Linn. Soc. 92, 211–219. 10.1111/j.1095-8312.2007.00820.x

52. Ohta, T., Gillespie, J.H., 1996. Development of neutral nearly neutral theories. Theor. Popul. Biol. 49, 128–142. 10.1006/tpbi.1996.0007

53. Okunowo, W., Ogunkanmi, A., 2010. Phytoremediation potential of some heavy metals by water hyacinth. Int. J. Biol. Chem. Sci. 4, 347–353. 10.4314/ijbcs.v4i2.58121

54. Otto, S.P., Whitton, J., 2000. Polyploid Incidence and Evolution. Annu. Rev. Genet. 34, 401–437. 10.1146/annurev.genet.34.1.401

55. Parisod, C., Holderegger, R., Brochmann, C., 2010. Evolutionary consequences of autopolyploidy. New Phytol. 186, 5–17. 10.1111/j.1469-8137.2009.03142.x

56. R core team, 2021. R: A language and environment for statistical computing.

57. Raabová, J., Fischer, M., Münzbergová, Z., 2008. Niche differentiation between diploid and hexaploid Aster amellus. Oecologia 158, 463–472. 10.1007/s00442-008-1156-1

58. Ramsey, J., 2007. Unreduced gametes and neopolyploids in natural populations of Achillea borealis (Asteraceae). Heredity 98, 143–150. 10.1038/sj.hdy.6800912

59. Ramsey, J., Schemske, D.W., 1998. Pathways, mechanisms, and rates of polyploid formation in flowering plants. Annu. Rev. Ecol. Syst. 29, 467–501. 10.1146/annurev.ecolsys.29.1.467

60. Rodríguez, D.J., 1996. A model for the establishment of polyploidy in plants. Am. Nat. 147, 33–46. 10.1086/285838

61. Segraves, K.A., Anneberg, T.J., 2016. Species interactions and plant polyploidy. Am. J. Bot. 103, 1326–1335. 10.3732/ajb.1500529

62. Shimizu, K.K., 2022. Robustness and the generalist niche of polyploid species: Genome shock or gradual evolution? Curr. Opin. Plant Biol. 69, 102292. 10.1016/j.pbi.2022.102292

63. Sonnleitner, M., Hülber, K., Flatscher, R., García, P.E., Winkler, M., Suda, J., Schönswetter, P., Schneeweiss, G.M., 2016. Ecological differentiation of diploid and polyploid cytotypes of Senecio carniolicus sensu lato (Asteraceae) is stronger in areas of sympatry. Ann. Bot. 117, 269–276. 10.1093/aob/mcv176

64. Stan Development Team, 2019. RStan: the R interface to Stan.

65. Stebbins, G.L., 1971. Chromosomal evolution in higher plants. Edward Arnold Ltd., London.

66. Stebbins, G.L., 1950. Variation and Evolution in Plants. Columbia University Press. 10.7312/steb94536

67. Sugiyama, S., 1998. Differentiation in competitive ability and cold tolerance between diploid and tetraploid cultivars in Lolium perenne. Euphytica 103, 55–59. 10.1023/A:1018322821118

68. Tayalé, A., Parisod, C., 2013. Natural Pathways to Polyploidy in Plants and Consequences for Genome Reorganization. Cytogenet. Genome Res. 140, 79–96. 10.1159/000351318

69. te Beest, M., Le Roux, J.J., Richardson, D.M., Brysting, A.K., Suda, J., Kubešová, M., Pyšek, P., 2012. The more the better? The role of polyploidy in facilitating plant invasions. Ann. Bot. 109, 19–45. 10.1093/aob/mcr277

70. Thompson, J.D., Lumaret, R., 1992. The evolutionary dynamics of polyploid plants: origins, establishment and persistence. Trends Ecol. Evol. 7, 302–307. 10.1016/0169-5347(92)90228-4

71. Thompson, K.A., Husband, B.C., Maherali, H., 2015. No influence of water limitation on the outcome of competition between diploid and tetraploid Chamerion angustifolium (Onagraceae). J. Ecol. 103, 733–741. 10.1111/1365-2745.12384

72. Van Drunen, W.E., Husband, B.C., 2018. Immediate vs. evolutionary consequences of polyploidy on clonal reproduction in an autopolyploid plant. Ann. Bot. 122, 195–205. 10.1093/aob/mcy071

73. Wickham, H., Averick, M., Bryan, J., Chang, W., McGowan, L., François, R., Grolemund, G., Hayes, A., Henry, L., Hester, J., Kuhn, M., Pedersen, T., Miller, E., Bache, S., Müller, K., Ooms, J., Robinson, D., Seidel, D., Spinu, V., Takahashi, K., Vaughan, D., Wilke, C., Woo, K., Yutani, H., 2019. Welcome to the Tidyverse. J. Open Source Softw. 4, 1686. 10.21105/joss.01686

74. Xu, S., Stapley, J., Gablenz, S., Boyer, J., Appenroth, K.J., Sree, K.S., Gershenzon, J., Widmer, A., Huber, M., 2019. Low genetic variation is associated with low mutation rate in the giant duckweed. Nat. Commun. 10, 8–13. 10.1038/s41467-019-09235-5

