## Appendix for "Polyploid - diploid coexistence in the greater duckweed *Spirodela polyrhiza*"

Invasion experiments test polyploid establishment and niche differentiation in greater duckweed Spirodela polyrhiza

### Appendix 1: Adapted Galbraith buffer

We used a nuclei extraction buffer containing 45mM MgCl2 + 6H2O, 30mM Sodium citrate + 2H2O, 20mM 4-morpholinepropane sulfonate (MOPS), 0.1% (vol/vol) Triton X-100 and 1% (mass/vol) Polyvinylpyrrolidone (PVP, Wu et al., 2023). This constitutes the buffer proposed by Galbraith and colleagues (1983), supplemented with PVP for binding cytosol phenolic compounds.

### Appendix 2: Statistical model description of tetraploid proportion models with calibration.

The proportion of tetraploid nuclei (${p4n}_{nuclei}$) is modelled as normally distributed (S1) and determined by the proportion of tetraploid individuals in the sample (${p4n}_{ind, i}$) with strain-specific absolute ($\alpha_{[strain]}$) and proportional offset ($\beta_{[strain]}$) on the logit scale (S2). The (logit of) proportion of tetraploid individuals in the sample (${p4n}_{ind, i}$) is, then, modelled as a normally distributed outcome (S3) determined by strain, salt-treatment and time (week). In the main text, we present models where we fixed all intercepts to the known initial tetraploid proportion of 0.05 (logit(0.05) = -2.94) for the tetraploid invasion experiment and 0.95 (logit(0.95) = 2.94) for the diploid invasion experiment. The model estimates the strain, salt-treatment and their interaction effect on the change in time of proportion: strain:week, salt:week and strain:salt:week (S4.1). We also estimate a variable slope in time for each replicated population (pop). We, furthermore, tested the same model with the difference that it estimated intercepts from the data (S4.2, supplementary materials, Appendix 5). In each case we modelled the tetraploid invasion experiment separately from the diploid invasion experiment.

$logit\left( {p4n}_{nuclei} \right) \sim Normal(\bar{p4n}_{nuclei}, \sigma_{nuclei})$	(S1)

$\bar{p4n}_{nuclei}= \alpha_{[strain]}+ \beta_{[strain]}* logit({p4n}_{ind, i})$	(S2)

$\alpha_{[strain]}, \beta_{[strain]} \sim Normal(0, 1)$	priors

$\sigma_{nuclei}\sim HalfCauchy\left( 0, 0.1 \right)$

$missing logit\left( {p4n}_{ind, i} \right) \sim Normal(\mu_{mi}, \sigma_{mi}$)

$\mu_{mi} \sim Normal(0, 2)$

$\sigma_{mi} \sim HalfCauchy($0, 1)

$logit({p4n}_{ind}) \sim Normal(\bar{p4n}_{ind}, \sigma_{ind})$	(S3)

$\mu_{ind} \sim0+{Intercept}_{exp}+ strain:week*salt:week+(0+week|pop)$	(S4.1)

${Intercept}_{tetraploid invasion}= -2.94$	priors

${Intercept}_{diploid invasion}=2.94$

$b \sim Normal\left( 0, 0.5 \right)$

$\sigma_{ind} \sim HalfCauchy(0.5, 0.5)$

${sd}_{pop} \sim HalfCauchy($0, 0.2)

${cor}_{pop} \sim LKJ(4)$

$\mu_{ind} \sim strain*salt*week+(1+week|pop)$	(S4.2)

${Intercept}_{tetraploid invasion} \sim Normal(-3, 1)$	priors

${Intercept}_{diploid invasion} \sim Normal(3, 1)$

$b \sim Normal\left( 0, 0.5 \right)$

$\sigma_{ind} \sim HalfCauchy(0.5, 0.5)$

${sd}_{pop} \sim HalfCauchy($0, 0.2)

${cor}_{pop} \sim LKJ(4)$

Priors were chosen to be informative but still wide enough to contain realistic parameter values. The prior b was used for all coefficients related to predictor variables in (S4.1) and (S4.2). We restricted the priors for $\sigma_{nuclei}$, $\sigma_{ind}$, and ${sd}_{pop}$ more severe than thought for the sake of model performance.

Models are fitted by first imputing the tetraploid individual proportion from the nuclei proportion and calibration data (S1, S2). This procedure applies a missing data imputation to the unknown (latent) individual proportion in the experiments that is estimated from what is found in the strain-specific calibration data (fig. S2). A missing data point is replaced by a prior that enables the estimation of each missing value. We proceed with 50 posterior draws of the imputed tetraploid individual proportion to retain the uncertainty in this imputation, and estimate the experimental effects across these 50 datasets.

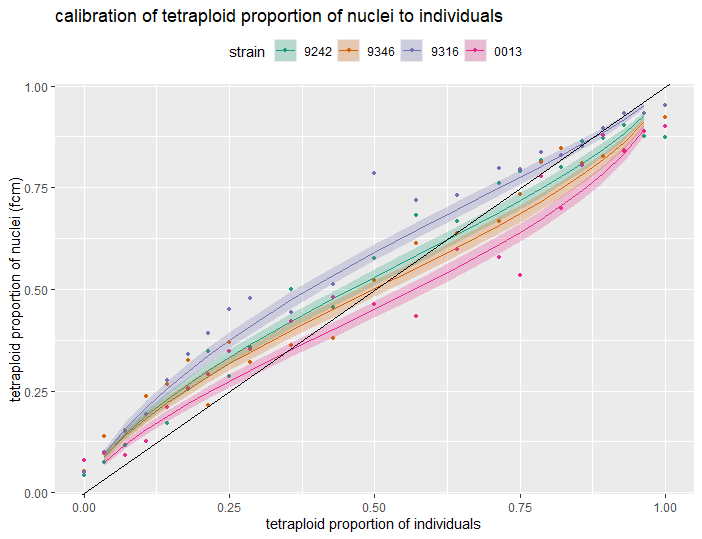

Fig. S2: the relation of tetraploid nuclei and individual proportion from the calibration data. We show a strain-specific mean calibration correction and the [0.09, 0.91] posterior likelihood interval. The black line indicates where individual proportion is equal to nuclei proportion.

### Appendix 3: Dry weight with measurement error from envelopes

Dried duckweed samples were weighed together with their envelope. We lacked a measurement of only plant dry weight, on which we wanted to infer the effects of strain, growth medium treatment and time but controlled for this by modelling the added measurement error. We estimated the added measurement error by weighing 30 separate envelopes that underwent the same drying and weighing procedure as the experimental dry weight measurements. Because we used larger envelopes in the first 4 weeks of the tetraploid invasion experiment, we also weighed 30 of those larger envelopes.

For both sizes of empty envelopes, we calculated the mean ($\bar{w}_{env\left[ size \right]}$) and standard deviation ($\sigma_{env\left[ size \right]}$) from 30 envelopes each. Empty envelopes underwent the same drying procedure as filled envelopes. We then used the difference between the total dry weight ($w_{total, i}$) and the mean envelope dry weight for that size ($\bar{w}_{env\left[ size \right]}$) as a response variable with a normal error distribution with standard deviation of the empty envelopes of that size (S5). The plant dry weight is modelled with a lognormal error distribution (S6) to reflect the strictly positive nature of plant weights and its median is determined by a logistic function of time (week) with three parameters (L: carrying capacity, k: growth rate and $x_{0}$: midpoint; S7) on which we model the effect of strain, growth medium treatment (salt), their interactions, and the effect of population to account for replicate-level differences in biomass and its growth (S8). The model can be formulated as follows:

$w_{total, i}- \bar{w}_{env\left[ size \right]} \sim Normal( w_{plant, i}, \sigma_{env\left[ size \right]})$	(S5)

$w_{plant, i} \sim Lognormal(\ln(\bar{w}_{plant}), \sigma_{plant})$	(S6)

$\bar{w}_{plant}= \frac{L}{1+e^{-k*(week-x_{0})}}$	(S7)

$L, k, x_{0}\sim strain*salt+(1|pop)$	(S8)

${Intercept}_{L, tetraploid invasion} \sim Normal\left( 5, 1 \right)$	priors

${Intercept}_{L,diploid invasion} \sim Normal\left( 4, 1 \right)$

${Intercept}_{k} \sim Normal\left( 0, 1 \right)$

${Intercept}_{x_{0}} \sim Normal\left( 0, 2 \right)$

$b_{L}, b_{k},b_{x_{0}} \sim Normal\left( 0, 0.5 \right)$

$\sigma_{env\left[ size \right]} \sim HalfCauchy\left( 0, 2 \right)$

${sd}_{L} \sim HalfCauchy\left( 0, 1 \right)$

${sd}_{k}, {sd}_{x_{0}} \sim HalfCauchy\left( 0, 0.1 \right)$

The total measured weight ($w_{total, i}$) is composed of the plant dry weight with its natural variation and the weight of a randomly sampled envelope with mean weight $\bar{w}_{env\left[ size \right]}$ and standard deviation $\sigma_{env\left[ size \right]}$. Because $w_{total, i}= w_{plant, i}+ Normal\left( \bar{w}_{env\left[ size \right]}, \sigma_{env\left[ size \right]} \right)= \bar{w}_{env\left[ size \right]}+ Normal\left( w_{plant, i}, \sigma_{env\left[ size \right]} \right)$, we can apply the easier-to-implement equation (S5).

Priors were chosen to be informative but still wide enough to contain realistic parameter values. The prior b was used for all coefficients related to predictor variables in (S8). The logistic relation is sensitive to different parameter values, which meant that we had to restrict some priors significantly in order to attain convergence on the model. We applied smaller prior distributions to the strain and salt effects. Because we measured a tenfold higher dry weight in the tetraploid compared to the diploid invasion experiment, we centered the prior for the intercept for carrying capacity close to the weights at the end of the experiment (or at least in proportion to what the sampled surface area represented compared to the total surface area of the box) for each experiment separately, but with wide enough distribution so that way higher carrying capacities were still sampled in the MCMC process. We also restricted priors for ${sd}_{k} \mathrm{and} {sd}_{x_{0}}$.

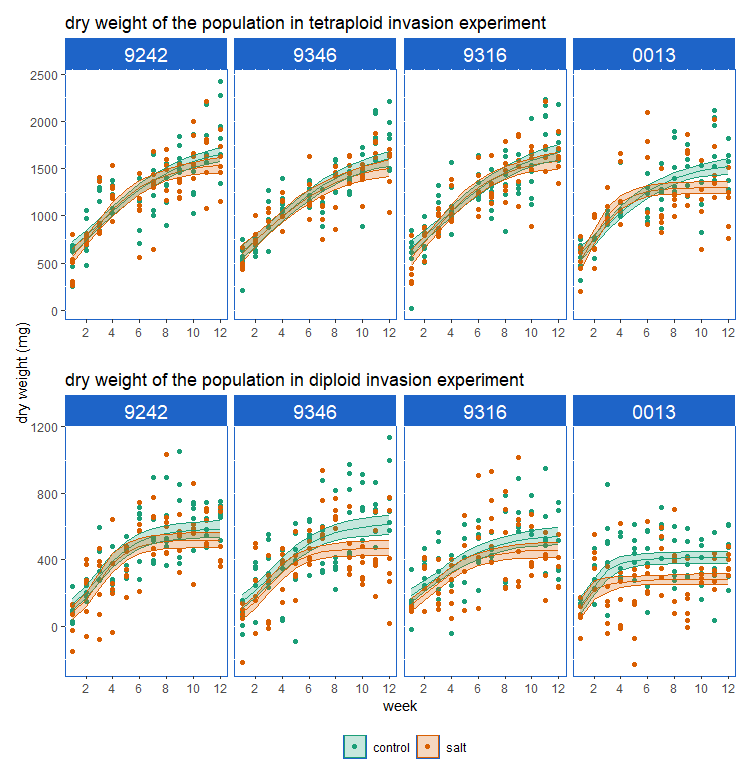

fig. S3.1: dry weight of the total population in the tetraploid invasion (upper) and diploid invasion (lower) experiment estimated on a weekly basis for all strains in control (green) and salt (orange) treatment. We fitted a logistic population growth model.

Total dry weight (i.e., of diploids and tetraploids combined) per replicate increased in both experiments. Biomass growth stagnated in all experimental combinations, showing the effects of competition in each microcosm. (fig. S3). Strains differed in the rate at which dry weight increased over the course of the experiment, but in all cases the rate of increase was equal or lower in the salt treatment than in the corresponding control treatment. Biomass growth also stagnated at a lower biomass in the salt treatment compared to the control. Tetraploid invasion experiments reached overall higher dry weights than the diploid invasion experiments. We used the most likely expected predicted (epred) posterior dry weight values to retrieve the most likely plant dry weights ($w_{plant, i}$) according to our model (Fig. S3.2). We used 10 draws of the posterior distribution to calculate cytotype population size.

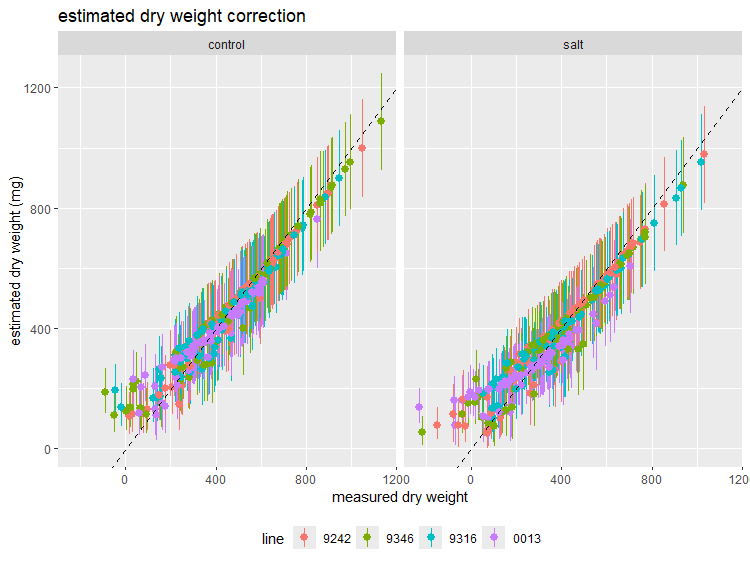

fig. S3.2: The estimated dry weight ($w_{plant, i}$) corrects the measured dry weight ($w_{total, i}- \bar{w}_{env\left[ size \right]}$) according to the measurement error that is expected from variation in the envelopes and in the expectation of positive plant dry weights. We plot the mean estimate (dot) and [0.04, 0.96] posterior likelihood interval. The positive plant weight expectation has the model mostly estimate that low measured dry weights have a likely real dry weight that is higher than the measured dry weight, as shown by the deviation from the dashed line.

### Appendix 4: count-to-weight conversion

We model the plant dry weight of a sample ($w_{i}$) as a normally distributed outcome determined by the number of duckweed fronds in a sample ($count, S10$) according to a conversion factor that is specific to strain-ploidy combination (c, S11).

$w_{i} \sim Normal(\bar{w}_{i}, \sigma)$	(S9)

$\bar{w}_{i}=c:count$	(S10)

$c= strain*ploidy$	(S11)

$b \sim Normal(0, 1)$	priors

$\sigma\sim HalfCauchy\left( 0, 1 \right)$

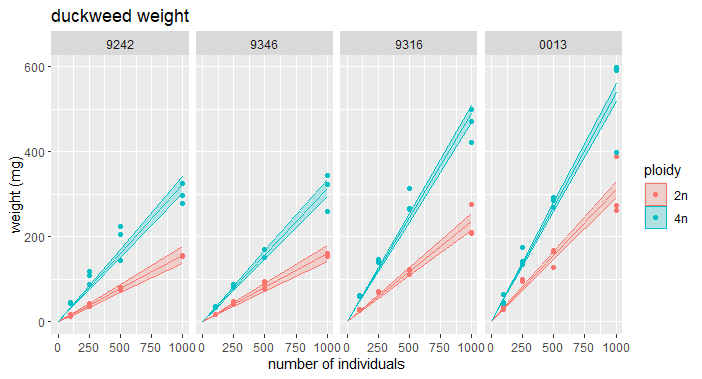

Fig S2.2: The estimated frond count to dry weight (mg) conversion. We plot the mean conversion and [0.09, 0.91] posterior likelihood interval.

### Appendix 5: Tetraploid proportion model with an estimated (unfixed) intercepts

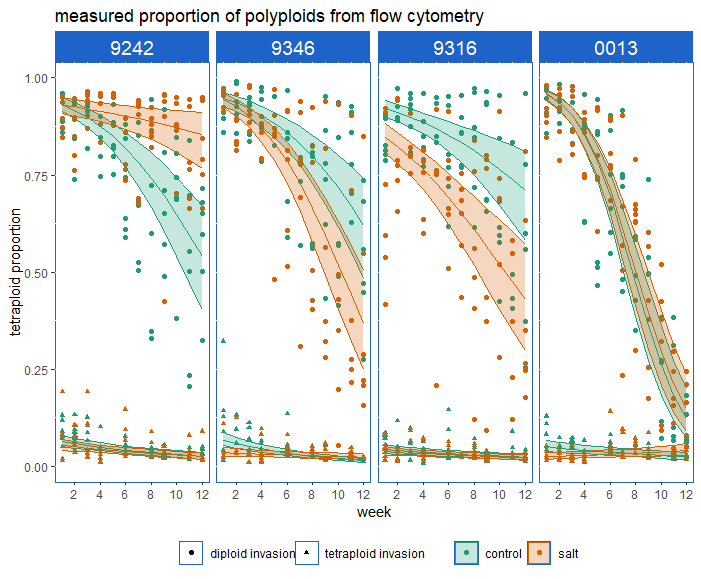

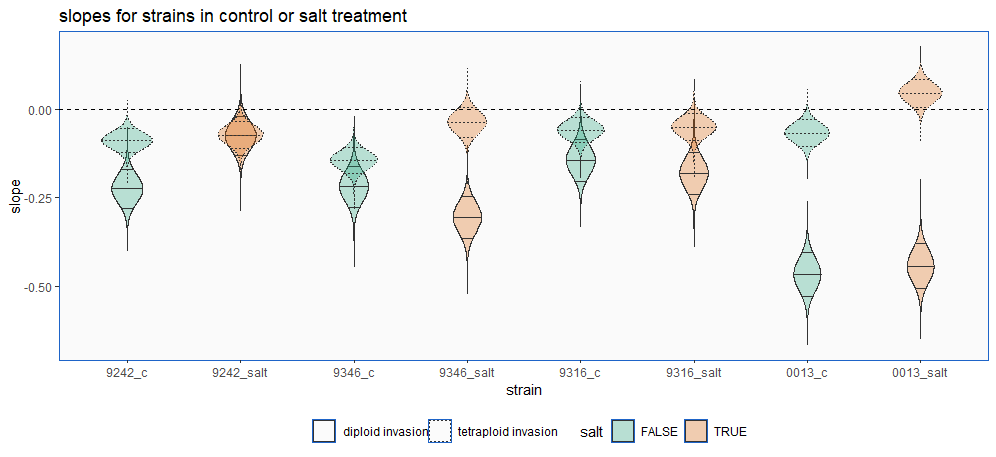

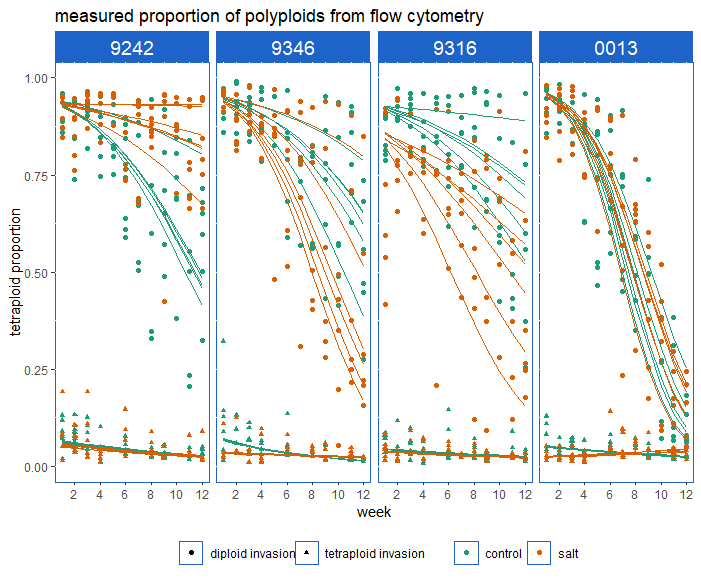

Fig. S5: Estimation of effects on tetraploid proportion when not fixing the intercept of the logistic regression model to the known initial tetraploid proportions. We plot posterior expected prediction, averaged over all replicates (upper), posterior slopes (middle) and posterior expected prediction per replicate population (lower) for all strains in the control (green) and salt (orange) environment. Not fixing the intercept reveals more or less the same trends compared to fixing them. These models estimate differences in intercept between some salt-treatment combination with sometimes consequences for the slope as well. The only qualitative difference with results that are discussed in the main text is that a lower estimated intercept of strain 0013 in salt in the tetraploid invasion experiment results in a positive slope in that combination. However, we are not convinced by this estimated increase in tetraploid proportion because of the inaccuracies of our method at lower proportions and the still relatively low estimated expected proportion (<0.1) at week 12.

### Appendix 6: tetraploid proportion expected from differences in exponential growth

If we assume that the cytotype’s populations grow exponentially, which is the case at low densities, we predict population size of cytotype i on day t ($N_{t, i}$) according to (S12). We determined the intrinsic growth rate from separate growth tests, for all four neotetraploid and four progenitor diploid strains in monocultures. They were performed in the same control (Hoagland) and salt (Hoagland + 2.5g/l NaCl) medium, in the same type of pots and otherwise same laboratory conditions as the invasion experiments. We tested 14-18 replicates in 3 batches, each started with 20 individuals (representatively sampled across all life stages) and counted the number of individuals after seven days. We calculated the strain-specific, per-week intrinsic growth rate of cytotype i ($r_{i}$) according to (S13, t = 1 week) and predict tetraploid proportion from only exponential growth ($p_{pred, t}$) according to (S14):

${N_{t, i}= N}_{0,i}* e^{r_{i}*t}$	(S12)

$r_{i}=\ln\left( N_{day 7, i} \right)-ln(N_{day 0, i})$	(S13)

$p_{pred, t}=\frac{N_{4n}}{N_{2n}+N_{4n}}= \frac{\frac{N_{4n}}{N_{2n}}}{1+ \frac{N_{4n}}{N_{2n}}}= \frac{\frac{N_{0,4n}}{N_{0,2n}}* \frac{e^{r_{4n}*t}}{e^{r_{2n}*t}}}{1+ \frac{N_{0,4n}}{N_{0,2n}}* \frac{e^{r_{4n}*t}}{e^{r_{2n}*t}}}= \frac{\frac{N_{0,4n}}{N_{0,2n}}* e^{{(r}_{4n}-r_{2n})*t}}{1+ \frac{N_{0,4n}}{N_{0,2n}}*e^{{(r}_{4n}-r_{2n})*t}}$ (S14)

With $N_{0, i}$the initial population size of cytotype i. Mean intrinsic growth rate for each cytotype and their difference are presented in table S5. We plotted these expectations, alongside the estimated tetraploid proportions in the main text (fig. 2, dashed lines).

$r \sim Normal(\bar{r}, \sigma)$	(S15)

$\bar{r}=Intercept+\beta_{\left[ strain X salt X ploidy \right]}+ \beta_{\left[ batch \right]}$	(S16)

$\beta_{\left[ batch \right]} \sim Normal(0, \sigma_{pop})$	(12)

$Intercept \sim Normal(1, 1)$	priors

$\beta_{\left[ strain X salt \right]} \sim Normal(0, 0.2)$

$\sigma_{pop} \sim HalfCauchy\left( 0, 0.2 \right)$

$\sigma\sim HalfCauchy\left( 0, 1 \right)$

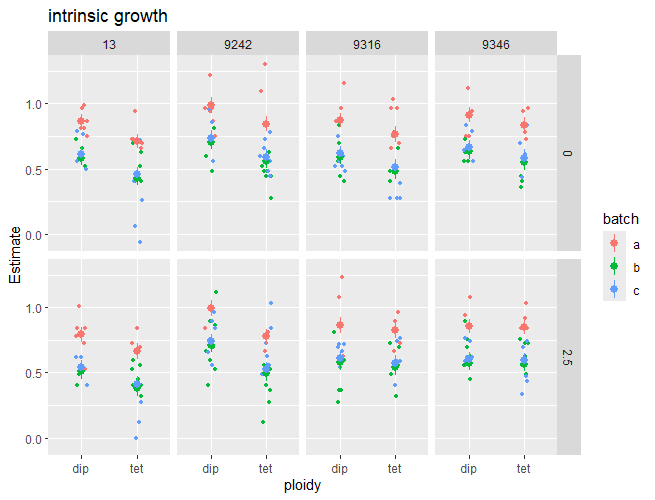

Fig S6: The measured (small) and estimated relative growth rate (RGR)of measurements from batch a (red) and b (blue) We plot the mean estimate and [0.09, 0.91] posterior likelihood interval.

|  | strain | salt | NaCl | batch | r_dip | r_tet | delta_r |
| --- | --- | --- | --- | --- | --- | --- | --- |
| 1 | 13 | 0 | 0 | a | 0.867072 | 0.712831 | -0.15424 |
| 2 | 13 | 1 | 2.5 | a | 0.79187 | 0.663918 | -0.12795 |
| 3 | 9242 | 0 | 0 | a | 0.988694 | 0.844717 | -0.14398 |
| 4 | 9242 | 1 | 2.5 | a | 0.994077 | 0.77631 | -0.21777 |
| 5 | 9316 | 0 | 0 | a | 0.872766 | 0.768382 | -0.10438 |
| 6 | 9316 | 1 | 2.5 | a | 0.863803 | 0.8273 | -0.0365 |
| 7 | 9346 | 0 | 0 | a | 0.915058 | 0.836885 | -0.07817 |
| 8 | 9346 | 1 | 2.5 | a | 0.856964 | 0.849114 | -0.00785 |
| 9 | 13 | 0 | 0 | b | 0.584023 | 0.429782 | -0.15424 |
| 10 | 13 | 1 | 2.5 | b | 0.508821 | 0.380869 | -0.12795 |
| 11 | 9242 | 0 | 0 | b | 0.705645 | 0.561668 | -0.14398 |
| 12 | 9242 | 1 | 2.5 | b | 0.711028 | 0.493262 | -0.21777 |
| 13 | 9316 | 0 | 0 | b | 0.589717 | 0.485333 | -0.10438 |
| 14 | 9316 | 1 | 2.5 | b | 0.580754 | 0.544251 | -0.0365 |
| 15 | 9346 | 0 | 0 | b | 0.63201 | 0.553836 | -0.07817 |
| 16 | 9346 | 1 | 2.5 | b | 0.573916 | 0.566066 | -0.00785 |
| 17 | 13 | 0 | 0 | c | 0.614317 | 0.460076 | -0.15424 |
| 18 | 13 | 1 | 2.5 | c | 0.539115 | 0.411163 | -0.12795 |
| 19 | 9242 | 0 | 0 | c | 0.735939 | 0.591962 | -0.14398 |
| 20 | 9242 | 1 | 2.5 | c | 0.741322 | 0.523556 | -0.21777 |
| 21 | 9316 | 0 | 0 | c | 0.620011 | 0.515627 | -0.10438 |
| 22 | 9316 | 1 | 2.5 | c | 0.611048 | 0.574545 | -0.0365 |
| 23 | 9346 | 0 | 0 | c | 0.662304 | 0.58413 | -0.07817 |
| 24 | 9346 | 1 | 2.5 | c | 0.604209 | 0.59636 | -0.00785 |

Table S6: Estimated mean intrinsic growth rates

We estimated a negative difference in relative growth rate (delta_r) for every strain treatment (NaCl) combination (fig. S6, table S6). The estimates for difference in relative growth rate (r_delta) are the same in each batch because we only estimate an effect of batch on the intercept of RGR, so modelling no interactions with ploidy. This means that we assume a constant difference across batches *a priori* (or estimate an average effect across the batches).

### Appendix 7: statistical model description of reciprocal differential equation model estimating population size of competing cytotypes.

We fit a statistical model for observed population size ($N_{i, t\_observed}$) of cytotype i at time t in the diploid invasion experiment. We calculated an estimated population size from the combined posterior distributions of the experimental cytotype proportion at time t, total weight at time t and the count-to-weight conversion. The observed population size is modelled as a lognormally distributed response with μ = $\log\left( N_{i, t} \right)$, which corresponds with a median at the expected population size ($N_{i, t}$, S17). Because population sizes are expected to be positive, we use a lognormal distribution. The starting population size of that cytotype ($N_{i, 0}$) is fixed at 10 and 190 for diploids and tetraploids respectively, in line with the experimental starting population sizes. The expected population size ($N_{i, t}$) is modelled as the starting population size added with the cumulative population growth until time t (continuous growth integrated from start until t; S18, S19). The continuous population growth is modelled with the reciprocal differential equations of Lotka-Volterra for competing species (Chesson, 2000; Godwin, Chang and Cardinale, 2020). $r_{i}$ represents the intrinsic growth rate of cytotype i, $\alpha_{i}$ the competition parameter of cytotype i on its own population growth and $\alpha_{i, j}$ the interaction parameter of cytotype j on the population growth of cytotype i. All six parameters of the population growth are modelled as the exponential of a sum of the intercept and coefficient for strain, for the interaction of strain and treatment (strain:salt) and a group-level effect per replicated population of the same strain-treatment combination (pop; S20, S21). We exponentiate the underlying effects to restrict intrinsic growth rates and competition parameters to be strictly positive, in line with the meaningful parameter space of this model.

$N_{i, t\_observed} \sim Lognormal(\log\left( N_{i, t} \right), \sigma)$	(S17)

$N_{2n, t}= N_{2n, 0}+ \int_{0}^{t} N_{2n, t}*r_{2n} (1- \alpha_{2n}N_{2n}- \alpha_{2n,4n}N_{4n})$ * dt	(S18)

$N_{4n, t}= N_{4n, 0}+ \int_{0}^{t} N_{4n, t}* r_{4n} (1- \alpha_{4n}N_{4n}- \alpha_{4n,2n}N_{2n})$ * dt	(S19)

$r_{i}=exp({Intercept}_{r}+ \beta_{\left[ strain \right]}+\beta_{\left[ strain:salt \right]}+ \beta_{\left[ pop \right]})$	(S20)

$\alpha_{i}, \alpha_{i, j}= exp({{Intercept}_{\alpha}+ \gamma}_{\left[ strain \right]}+\gamma_{\left[ strain:salt \right]}+ \gamma_{\left[ pop \right]})$	(S21)

$\sigma\sim Exponential(0.5)$	priors

$N_{2n, 0}= 10$

$N_{4n, 0}= 190$

$\beta_{\left[ pop \right]}, \gamma_{\left[ pop \right]} \sim Normal(0, \sigma_{\left[ pop \right]})$

${Intercept}_{r}, \beta\sim Normal(0, 1)$

$\gamma\sim Normal(0, 0.5)$

${Intercept}_{\alpha} \sim Normal(-7, 0.5)$

$\sigma_{\left[ pop \right]} \sim HalfCauchy(0, 0.1)$

Priors were chosen to be informative but still wide enough to contain realistic parameter values. We choose a prior for the intercept of each competition parameter centered around -7, because of the low expected value of these parameters with population size well above 10³. Because of the exponentiated effects, a prior centered around -7 represents competition parameters sampled around e^-7^ ≈10^-3^. We also had to restrict the standard deviation of the group-level effect ($\sigma_{\left[ pop \right]}$) for model performance.

We show the estimated a posterior distribution of average population growth parameters across all population replicates according to table S7. We also calculated the difference in intrinsic growth rate (delta_r) for each strain-treatment combination to compare to the calculated difference from intrinsic growth rate tests (table S6). The model estimated differences in the same range (most between -0.2 and 0) but estimates an extremely large difference for 9316 in salt, which was measured as the second smallest difference in the growth tests. This discrepancy between model estimation and intrinsic growth tests may indicate inaccuracies in the intrinsic growth parameter from experimental data (propagated inaccuracies) or a discrepancy in intrinsic growth tests.

| line | salt | r4n | r2n | a4n | a4n, 2n | a2n, 4n | a2n | delta_r |
| --- | --- | --- | --- | --- | --- | --- | --- | --- |
| 9242 | 0 | 0.78176 | 0.905814 | 0.000596 | 0.000469 | 0.000357 | 0.00061 | -0.12405 |
| 9242 | 1 | 0.5718 | 0.610424 | 0.000544 | 0.000426 | 0.000432 | 0.000559 | -0.03862 |
| 9346 | 0 | 0.730024 | 0.812556 | 0.000569 | 0.000323 | 0.000304 | 0.000541 | -0.08253 |
| 9346 | 1 | 0.629165 | 0.817689 | 0.0007 | 0.000509 | 0.000354 | 0.000447 | -0.18852 |
| 9316 | 0 | 0.922064 | 0.929404 | 0.001042 | 0.000572 | 0.000781 | 0.000628 | -0.00734 |
| 9316 | 1 | 0.585588 | 1.216859 | 0.001165 | 0.000675 | 0.000961 | 0.000687 | -0.63127 |
| 0013 | 0 | 0.759955 | 0.948477 | 0.001113 | 0.001577 | 0.000445 | 0.000913 | -0.18852 |
| 0013 | 1 | 0.702367 | 0.801928 | 0.001748 | 0.00219 | 0.000595 | 0.00115 | -0.09956 |

Table S7: Estimated average intrinsic growth (r) and competition parameter (a) for diploids (2n) and tetraploids (4n) across all replicated populations.

### Appendix 8: per-replicate tetraploid proportion

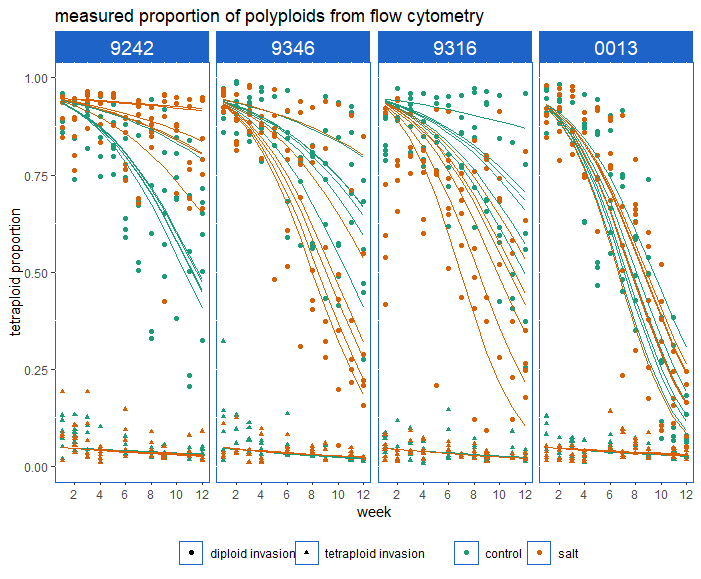

Fig. S8: tetraploid proportion trajectory for each replicated invasion experiment. Top: diploid invasion, bottom: tetraploid invasion.

Replicates of the same strain-treatment combination varied considerably in the diploid invasion experiments (fig. S8). For instance, one replicate shows a stable tetraploid proportion for strain 9242 in salt while others clearly declined. Earlier random variation in population size compounds towards the end of our experiment due to the multiplicative nature of population growth. This estimated variance between replicates was not caused by fixing the intercept (fig. S6).We also note the substantial variation within each replicate, unexplained by a steady change in tetraploid proportion. Combined with the challenge to make accurate proportion estimations from mixed cytotype flow cytometry, we introduced a larger sampling error by sampling 50 individuals that were often connected in groups of two, three or four instead of 50 independent individuals.

### References

Chesson, P. (2000) ‘Mechanisms of Maintenance of Species Diversity’, *Annual Review of Ecology and Systematics*, 31(1), pp. 343–366. Available at: https://doi.org/10.1146/annurev.ecolsys.31.1.343.

Galbraith, D.W. *et al.* (1983) ‘Rapid flow cytometric analysis of the cell cycle in intact plant tissues’, *Science (New York, N.Y.)*, 220(4601), pp. 1049–1051. Available at: https://doi.org/10.1126/science.220.4601.1049.

Godwin, C.M., Chang, F.-H. and Cardinale, B.J. (2020) ‘An empiricist’s guide to modern coexistence theory for competitive communities’, *Oikos*, 129(8), pp. 1109–1127. Available at: https://doi.org/10.1111/oik.06957.

Wu, T. *et al.* (2023) ‘Studying Whole-Genome Duplication Using Experimental Evolution of Spirodela polyrhiza’, in Y. Van de Peer (ed.) *Polyploidy: Methods and Protocols*. New York, NY: Springer US, pp. 373–390. Available at: https://doi.org/10.1007/978-1-0716-2561-3_19.
